# Cooperation between bacteriocytes and endosymbionts drives function and development of symbiotic cells in mussel holobionts

**DOI:** 10.1101/2023.08.18.553804

**Authors:** Hao Chen, Mengna Li, Zhaoshan Zhong, Minxiao Wang, Chao Lian, Guanghui Han, Hao Wang, Li Zhou, Huan Zhang, Lei Cao, Chaolun Li

**Author notes:** Correspondence and requests for materials should be addressed to H.C., M.W. or C.L. These authors contributed equally to this work (Hao Chen, Mengna Li, Zhaoshan Zhong).

## Abstract

Symbiosis drives the adaptation and evolution in multicellular organisms. Modeling the function and development of symbiotic cells/organs in holobionts is yet challenging. Here, we surveyed the molecular function and developmental trajectory of bacteriocyte lineage in non-model deep-sea mussels by constructing a high-resolution single-cell expression atlas of gill tissue. We show that mussel bacteriocytes optimized immune processes to facilitate recognition, engulfment, and elimination of endosymbionts, and interacted with them intimately in sterol, carbohydrate, and ammonia metabolism. Additionally, the bacteriocytes could arise from three different stem cells as well as bacteriocytes themselves. In particular, we showed that the molecular functions and developmental process of bacteriocytes were guided by the same set of regulatory networks and dynamically altered regarding to symbiont abundance via sterol-related signaling. The coordination in the functions and development of bacteriocytes and between the host and symbionts underlies the interdependency of symbiosis, and drives the deep-sea adaptation of mussels.

## Introduction

Symbiosis is a major force in the acquisition of novel adaptive traits, expansion of ecologic range, and shaping the biodiversity and evolution of eukaryotic organisms ^1^. To establish symbiotic associations, majority of animal hosts have formed highly complex symbiotic cells and organs (such as bacteriocytes of aphid, light organ of bobtail squid, trophosome of tubeworm), which harbor the symbionts (especially the endosymbionts) and provide mutualist relationships. Knowledge on the function and development of symbiotic cells and organs are therefore necessary to understand the formation and evolution of symbiosis ^2^. Along with a growing number of studies, the function of, and symbiotic interactions within, symbiotic cells and organs are becoming clear in some model holobionts ^3–5^. However, characterizing the function and development of symbiotic cells and organs remains challenging in non-model holobionts ^6^. Furthermore, questions on whether symbionts influence the function and development of symbiotic cells, and the possible mechanisms involved in this activity, remain debated and may vary greatly across species ^7–9^.

Since their first discovery in 1977, the chemosynthetic ecosystems in deep-sea cold seeps and hydrothermal vents have attracted much attention ^10^. Noticeably, majority of the endemic invertebrates in these ecosystems have formed closed symbiotic association with the chemosynthetic bacteria to obtain nutrition ^11^. Among them, the mollusks (especially bivalves from the *Mytilidae, Vesicomyidae, Solemyidae, Thyasiridae,* and *Lucinidae* families) are particular interesting as the chemosymbiosis in mollusks can vary greatly depending on the symbiont location (extracellular or intracellular), symbiont type (methanotroph, thiotroph, or both), and transmission mode (vertical or horizontal) ^12^. Moreover, although certain mollusks have developed bacteriocytes in their gill tissue to house chemosynthetic endosymbionts, others may lose their symbionts in particular environments and experience a “reverse” evolution akin to their non-symbiotic shallow water counterparts ^13,14^. For these reasons, the mollusks are therefore regarded as a promising model in investigating the chemosymbiosis, and especially the function and development of symbiotic cells and organs.

Among the deep-sea mollusks, the *Bathymodiolinae* mussels are known to harbor endosymbiotic methanotrophs and/or thiotrophs in their specialized gill epithelial cells (bacteriocytes) ^15,16^. It is estimated that adult deep-sea mussels contain 10^8^ to 10^9^ bacteriocytes with hundreds of methanotrophic endosymbionts or thousands of thiotrophic endosymbionts harbored inside a single bacteriocyte ^17^. Recently, several pathways and genes potentially involved in the metabolic and immune interaction between mussel and endosymbionts are proposed, highlighting the unique way of host-symbiont interaction ^18–22^. Additionally, it is also demonstrated that new bacteriocytes could arise from gill cells adjacent to growth zones, in which symbionts were colonized from ontogenetically older bacteriocytes ^23^. A more insightful finding is that the structure of new bacteriocytes would change drastically after symbiont infection, indicating the potential role of symbionts in the function and development of bacteriocytes ^23,24^. However, due to the extensive heterogeneity of gill tissue and limits in the isolation and *in vitro* culture of bacteriocytes and endosymbionts, the exact function and development process of bacteriocytes, as well as the mechanisms beneath these processes, still remain obscure and debatable. With recent advances in single cell transcriptome and spatial transcriptome technologies, it is now possible to reveal the function and development of symbiotic cells and organs in either model or non-model organisms, and with and without host cell lineages and culturable symbionts ^25–27^. Recently, we have constructed a comprehensive cell atlas of the gill tissue in the methanotrophic deep-sea mussel *Gigantidas platifrons* using single-nucleus RNA sequencing ^28^. The successful identification of three stem cell lineages and bacteriocyte lineage provides opportunity to reveal the function and development of bacteriocytes, and to survey how symbionts influence these processes. With help of an updated high-resolution single-cell transcriptome and spatial transcriptome, we here conducted a comprehensive analysis by using the phagocytosis assay, 5-ethynyl 2’-deoxyuridine (EdU) labeling assay, and 3D electron microscopy data to characterize the molecular function and development trajectory of bacteriocytes. We also conducted an *in situ* decolonization assay to compare our data with those obtained on the decolonized mussel and have successfully demonstrated the potential influence of symbionts on the function and development of symbiotic cells.

## Results

### Functional landscape of bacteriocyte lineage

To improve the single cell transcriptome performance and characterize the expression pattern of key cell lineages such as bacteriocytes and stem cells, we here employed two groups of deep-sea mussels in the present study and conducted both single-cell transcriptome (scRNA-seq, Chromium platform of 10x Genomics) and spatial transcriptomics (ST-seq, Visium platform of 10x Genomics) analysis (Fig. 1A). In particular, mussels collected from the seepage region (methane concentration up to 31,227 ppm, designated as the InS group, n=14) were used to represent normal mussels with a fully symbiotic state; mussels transplanted *in situ* to a low methane region (methane concentration about 800 ppm) for 604 days with an average 76.5% decrease of endosymbionts were used to represent partially decolonized mussels (DeC group, n=5). Principal co-ordinates analysis (PCoA) based on the meta-transcriptome data of the two groups showed robust expressional stability within individuals of the same group and strong variations between individuals of different groups (Supplementary Fig. S1A–B). Two individual mussels with similar body size were randomly selected from each group and subjected for scRNA-seq and ST-seq. To include more possible cell types, the dorsal–middle region of the gill (containing both descending and ascending gill filaments and adjacent to posterior end of mussel) was employed to scRNA-seq after cell nuclei extraction. In addition, the cross-sectioned middle region of the gill (dorsal view) was also used for ST-seq to verify the identified cell types (Fig. 1A). As a result, approximately 26,707 cells with 37,585 mean reads per cell, and over 3,600 barcoded spots with 172,705 mean reads per spot, were obtained for the two samples evaluated using scRNA-seq and ST-seq respectively (Supplementary Table 1). Transcript coverage profiling indicated that sequencing depth of all samples was saturated for scRNA-seq and ST-seq (Supplementary Fig. S1C–F), ensuring the reliability of subsequent analyses.

**Fig. 1.**
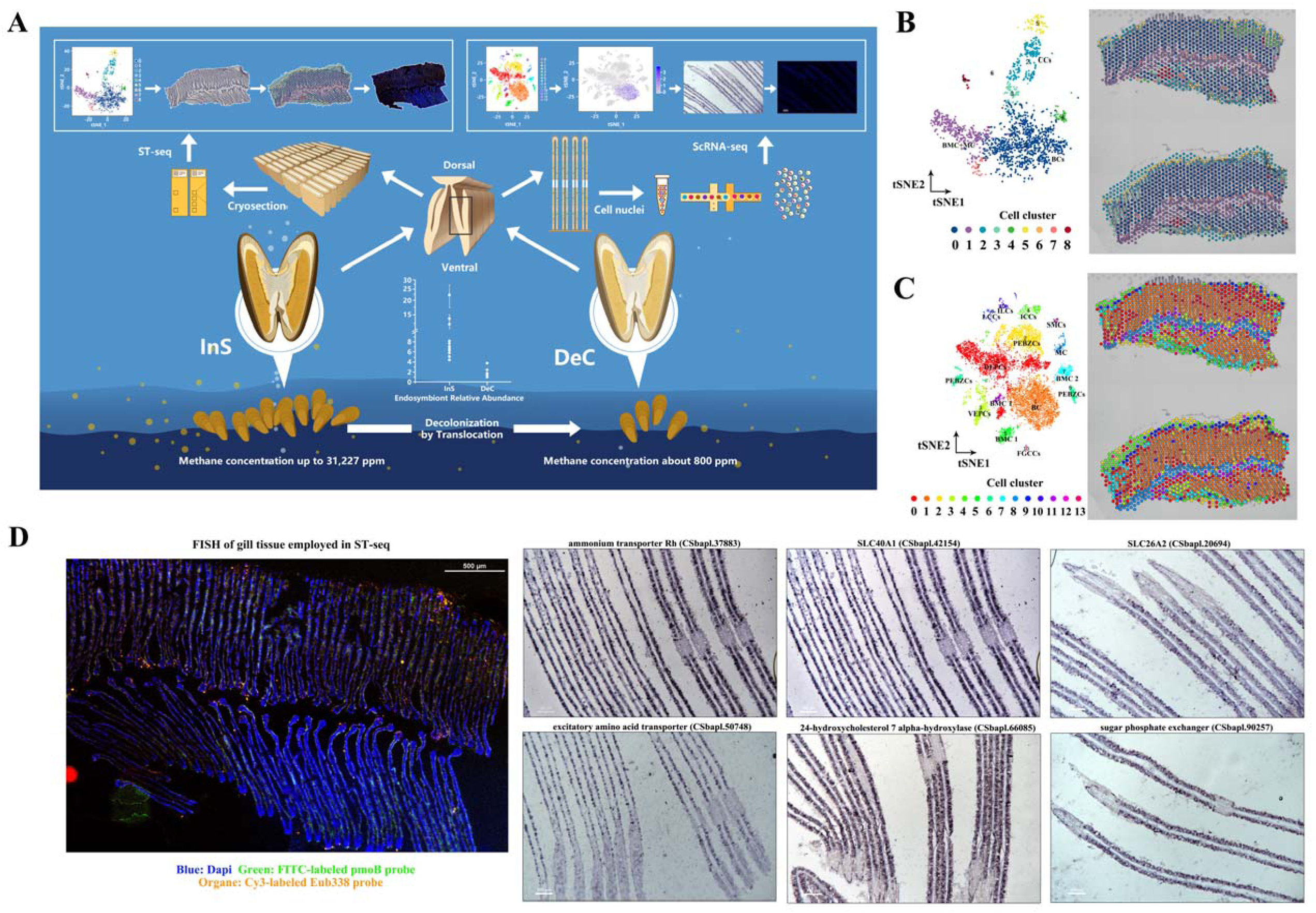
Spatially resolved single-cell transcriptomic atlas of deep-sea mussel gill. (A) Experimental workflow for, and analysis of, single-cell transcriptome (scRNA-seq) and spatial transcriptome (ST-seq) of gill tissue in fully symbiotic (InS group) and partially decolonized (DeC) deep-sea mussels. A marked decrease of the endosymbiont is observed after *in situ* translocation assay (n=14 for the InS group, n=5 for the DeC group). (B) t-SNE projection of spatial transcriptome clustered by gene expression in the InS group, with color assigned by cell type (left). Projection of cell clusters onto spatial transcriptome barcoded spots (right) in the InS group. Two successive sections were used in the same capture region of spatial transcriptome as technique replicates. (C) t-SNE projection of single-cell transcriptome clustered by gene expression in the InS group, with color assigned by cell type (left). Spatial transcriptome barcoded spots labeled using scRNA-seq cell type with maximum prediction score (right). (D) Fluorescent *in situ hybridization* (FISH) of endosymbionts with successive gill sections for ST-seq (InS group) and ISH of bacteriocytes marker genes.

Consequently, a total of 14 cell clusters were identified here based on our scRNA-seq data, among which 13 clusters were found sharing majority of marker genes identified in our previous study ^28^. For example, the bacteriocytes were found transcribed a total of 199 candidate marker genes (with expression levels at least 1.5-fold higher than that of other cell cluster; *p*-value less than 0.01, Supplementary Table 2) based on scRNA-seq data of both the InS and DeC groups, which consisted 10 out of 25 candidate marker genes that identified previously ^28^. To further verify the identified cell types, we combined scRNA-seq- and ST-seq-derived data using Seurat-v3.2 anchor-based integration and projecting all identified cell clusters from the scRNA-seq data onto gill tissue data of ST-seq (Fig. 1B–C). In support of the scRNA-seq data, all annotated bacteriocytes in the scRNA-seq were found co-located with MOB signals of the gill filament (Fig. 1D) while majority of the annotated marker genes (83/199) were also found abundantly expressed in ST-seq data of InS group. Given to the shared marker genes, we have successfully identified the bacteriocytes (cluster 1), three types of stem cells [dorsal end proliferation cells (DEPCs, cluster 0), posterior end budding zone cells (PEBZCs, cluster 2 and 6)], five types of ciliary and smooth muscle cells [intercalary cells (ICCs, cluster 4), lateral ciliary cells (LCCs, cluster 9), food grove ciliary cells (FGCCs, cluster 13), and smooth muscle cells (SMCs, cluster 12)], five supportive cells [basal membrane cell 1 (BMC1, cluster 5 and 11), BMC2 (cluster 7), mucus cell (cluster 8), and inter lamina cells (ILCs, cluster 10)] in our scRNA-seq data.

In addition, the median number of identified genes per cell and total number of candidate marker genes (Supplementary Table 1, 2) have been improved substantially in comparison with our previous study (with only 631 identified genes per cell per sample, and 25 marker genes for bacteriocytes, 15 marker genes for stem-like cells) ^28^, which reassures the characterization of function and development of bacteriocytes. Indeed, the improved scRNA-seq performance provided more detailed information on the biological processes of bacteriocytes. For example, function enrichment analysis of marker genes (Supplementary Fig. S2A) showed that the bacteriocytes largely encoded genes involved in metabolic processes (such as biosynthesis and transport of carbohydrates, lipids, amino acids, and vitamins) and symbiosis-related immune responses (including GO: 0002376, GO: 0009617, GO: 0044111, and GO: 0044403) (Supplementary Fig. S2). Particularly, we noticed that a total of 67 out of 199 marker genes, such as excitatory amino acid transporter 1 (*SLC1A3*), monocarboxylate transporter 13 (*SLC16A14*), cathepsin-L (*CTL*), acid phosphatase type 7 (*ACP7*), and myoneurin (*MYNN*), were consistently expressed by bacteriocytes in the InS and DeC groups regardless of fluctuations in symbiont abundance. Comparatively, the rest of the 132 out of 199 marker genes, including solute carrier family 40 member 1 (*SLC40A1*), ammonium transporter Rh type A (*RHBG-A*), solute carrier family 26 member 10 (*SLC26A2*), 24-hydroxycholesterol 7 alpha-hydroxylase (*CYP39A1*, EC1.14.14.26), sugar phosphate exchanger 2 (*SLC37A2*), zinc finger protein 271 (*ZNF271*), and ETS-related transcription factor (*EHF*) were downregulated in the DeC group in response to symbiont decolonization (Supplementary Table 3).

### Symbiosis-related immune processes in bacteriocytes

With merely the innate immunity, how invertebrate host forged the symbiotic association with the specific chemosynthetic bacteria remains intriguing. We therefore screened for the possible genes and pathways that participate in the establishment and maintenance of symbiosis in bacteriocyte lineages. Although PRRs, such as TLRs and PGRPs, could play vital roles in the recognition of symbionts in other holobionts ^29^, we found that only several pattern recognition receptors (PRRs) were highly expressed in bacteriocytes of fully-symbiotic mussels (InS group) or decolonized mussels (DeC group), while numerous PRRs showed a relatively low level of expression or even failed to be characterized (e.g., approximately 55 out of 146 TLRs were not characterized by scRNA-seq). Nevertheless, one *TLR2* gene were annotated as markers for bacteriocytes, respectively (Fig. 2A, Supplementary Table 2). In addition, two PRRs (*rhamnose-binding lectin* and *TLR2*) showed increased expression levels in bacteriocytes after decolonization (Supplementary Table 3).

**Fig. 2.**
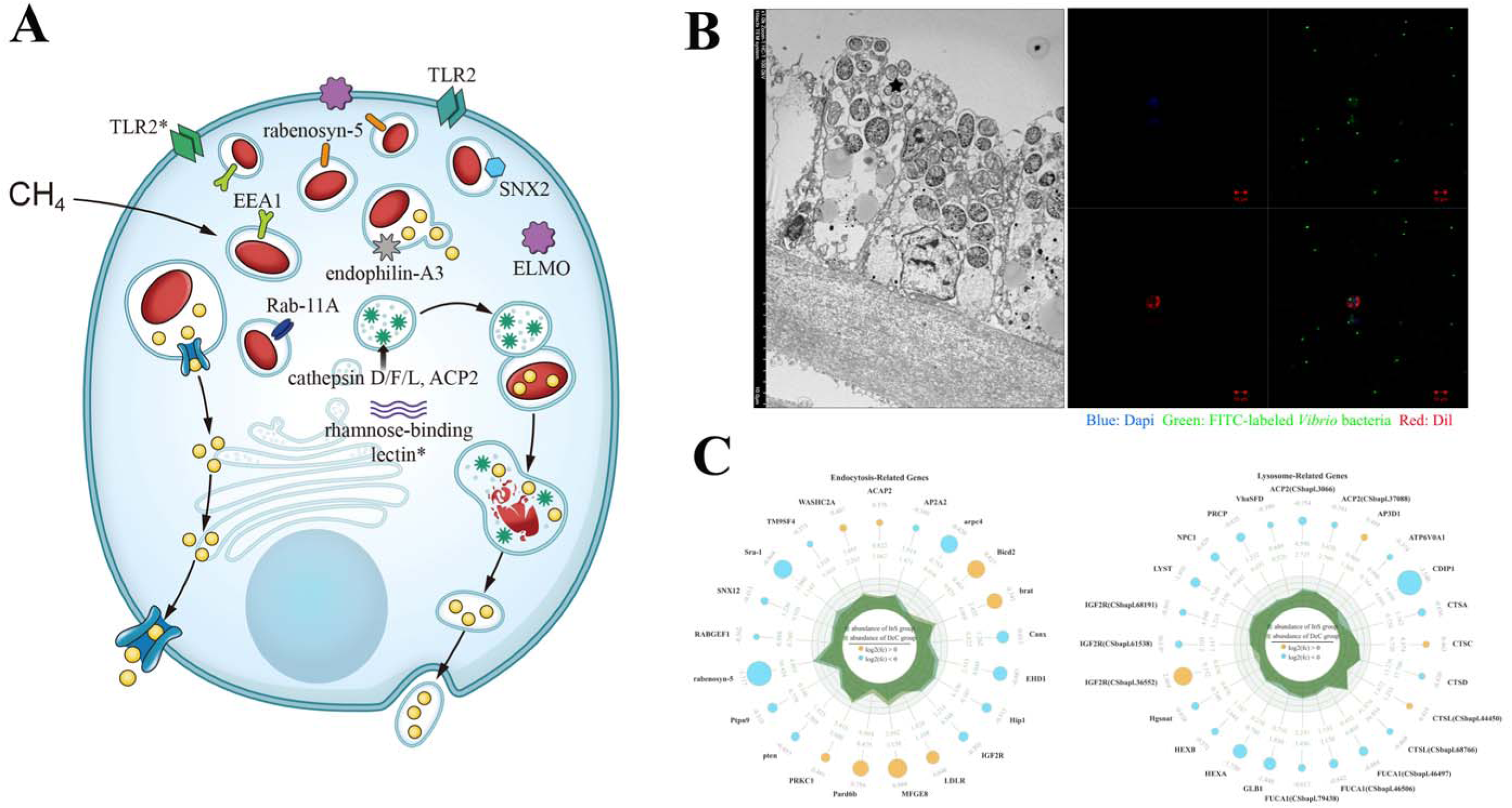
Immune genes and processes involved in symbiosis. (A) Schematic drawing of immune-related marker genes of bacteriocytes. (B) Engulfment of symbionts (labeled with star) by bacteriocytes as revealed by TEM and phagocytosis of FITC-labeled *Vibrio alginolyticus* by bacteriocytes *in vitro*. (C) Radar chart showing expression pattern of endocytosis- and lysosome-related genes of bacteriocytes in the InS and DeC groups. The size of blue/yellow colored apical circles represents the absolute value of log_2_(fold changes), where blue indicates downregulation, and yellow indicates upregulation, in partially decolonized mussels. Green and dark green colored inner diagram represents the abundance of a given gene in decolonized and fully symbiotic mussels, respectively.

At the meanwhile, we have detected abundant expression of endocytosis- and lysosome-related marker genes in bacteriocytes of fully-symbiotic mussels (Fig. 2A, Supplementary Table 2). The coordination among these genes could facilitate phagocytosis and lysosome-mediated digestion of bacteria, which are often observed in bacteriocytes and play crucial in establishment and maintenance of symbiosis (Fig. 2B, Supplementary Fig. S2B). Noticeably, we noted that majority of endocytosis- and lysosome-related genes were down-regulated when symbionts decreased (Fig. 2C, Supplementary Table 3), implying that endocytosis and lysosome-mediated digestion could be vigorously regulated by the host depending on symbiont abundance. In addition, although massive lysosome-related genes were abundantly expressed in bacteriocytes while some endosymbionts were being digested by lysosomes, our EdU labeling assay and 3D electron microscopy analysis yet showed that few endosymbionts (3 out of 408 symbionts for an individual bacteriocyte) were proliferating inside normal bacteriocytes of adult mussels and even fewer were being captured from environment (InS group, Supplementary Fig. S2C, Supplementary Video.1).

### Metabolic interaction between the host and symbionts

As the interface of symbiotic association, the deep-sea mussels are known obtain and transport symbionts-deprived nutrients in bacteriocytes. As observed in the function enrichment analysis, metabolic genes and processes were significantly up-regulated in bacteriocytes than other cells, highlighting the unique role of bacteriocytes. Noticeably, majority of metabolic marker genes were involved in biosynthesis and transport of carbohydrates, lipids, amino acids, and vitamins, implying metabolic interactions between the deep-sea mussels and symbionts. Among these marker genes, the *CYP39A1* (EC1.14.14.26) gene was found massively transcribed in bacteriocytes of fully symbiotic mussels (InS group) based on both scRNA-seq and ST-seq data (Fig. 3A). As the key enzyme in the turnover of sterol intermediates such as presqualene-PP, squalene, (S)-squalene-2,3-epoxide, and 4,4-dimethyl-cholesta-8,14,24-trienol, the unique expression pattern of *CYP39A1* (EC1.14.14.26) gene inside bacteriocytes was further confirmed by FISH and IHC assay. However, we also noticed that the mussel hosts are lack of related enzymes such as EC2.5.1.21, EC1.14.1417, EC1.14.14154, and EC1.14.1536 to synthesize the above sterol intermediates. Consistently, meta-transcriptome data showed that methanotrophic endosymbionts were robustly transcribing enzymes such as EC2.5.1.21 and EC1.14.14154 under normal condition (the top 10% highly expressed genes in the InS group) in a complementary manner with the host (Fig. 3A, Supplementary Fig. S3A–D, Supplementary Table 4), highlighting the metabolic dependency of mussel hosts on sterol intermediates of symbionts ^30^.

**Fig. 3.**
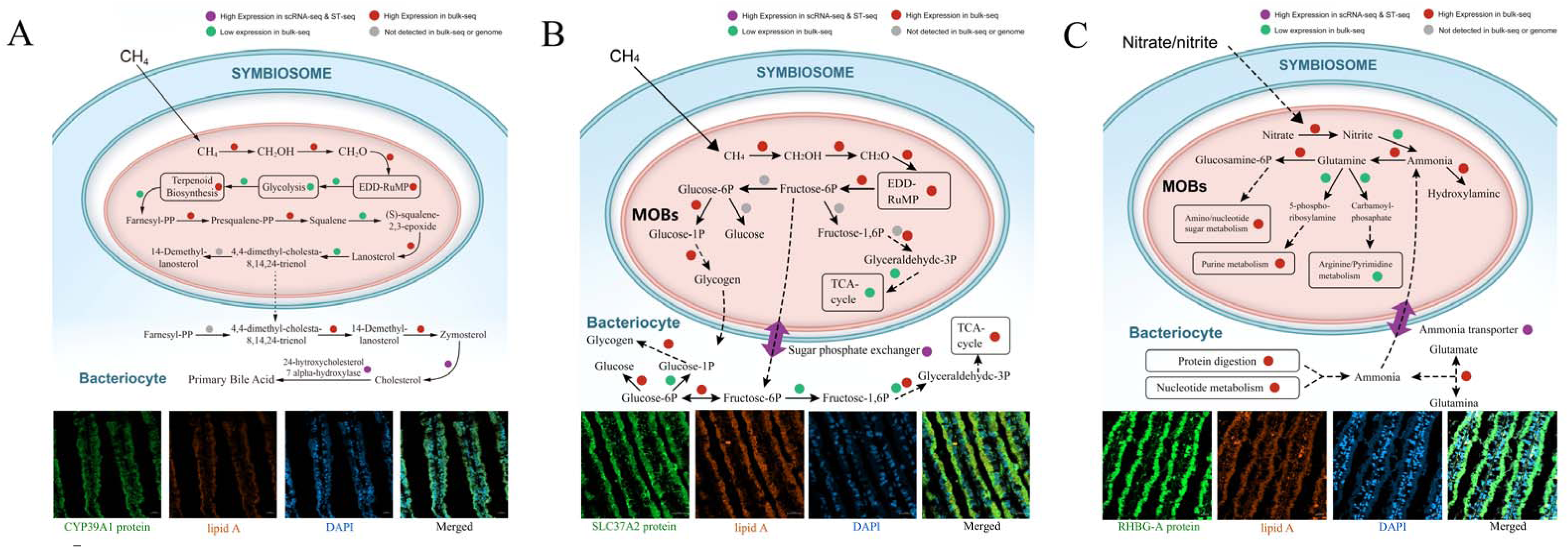
Metabolic interactions between mussel host and endosymbionts. (A) ScRNA/ST-seq and meta-transcriptome data show an intimate interaction of sterol metabolism between the host and symbionts. In support of this, immunofluorescence (IF) assay (using gills of the InS group) of the 24-hydroxycholesterol 7 alpha-hydroxylase (CYP39A1) protein shows that CYP39A1 proteins are widely distributed across bacteriocytes. In addition, a more intensive signal of CYP39A1 protein could also be observed at the apical region that enriched with endosymbionts (indicated by lipid A signals). (B) ScRNA/ST-seq and meta-transcriptome data show intimate interaction of glucose/glycogen metabolism between the host and symbionts. In support of this, IF assay of the sugar phosphate exchanger (SLC37A2) protein shows the co-location of SLC37A2 proteins with endosymbionts inside bacteriocytes. (C) ScRNA/ST-seq and meta-transcriptome data show intimate interactions of ammonia metabolism between the host and symbionts. In support of this, IF assay of the ammonium transporter Rh (RHBG-A) protein shows co-location of RHBG-A proteins with endosymbionts inside bacteriocytes.

It is also noticed that both host and endosymbionts were massively transcribing key genes involved in gluconeogenesis and glycogen biosynthesis (including EC2.4.1.18, 2.4.1.21, 2.7.7.27, and 5.4.2.2) as demonstrated by meta-transcriptome data (the top 10% of highly expressed genes in the InS group, Fig. 3B, Supplementary Fig. S4, Supplementary Fig. S5, Supplementary Table 4). While these results implied a robust production of glycogen in mussels and endosymbionts, we also noted that multiple enzymes in the gluconeogenesis pathway of endosymbiont were either insufficiently expressed (EC5.3.1.9) or completely absent from the genome (EC3.1.3.11, EC2.7.1.11, and EC4.1.2.13), which could result in overproduction of fructose-6P and shortages of glucose-6P, fructose-1,6P_2_, glyceraldehyde-3P, and glucose, and therefore slow down the production of glycogen in symbionts and host. Conversely, we found that all of these enzymes were transcribed by bacteriocytes, while a sugar phosphate exchanger gene *SLC37A2* (responsible for transmembrane transport of fructose-6P, glucose-6P, fructose-1,6P_2_, and glyceraldehyde-3P) were abundantly expressed by bacteriocytes even under normal conditions (scRNA-seq and ST-seq of InS group) with proteins co-localizing with endosymbionts (Fig. 3B, Supplementary Fig. S5). Complementation in carbohydrate metabolism strongly suggested that symbionts could supply fructose-6P directly to the host in exchange for gluconeogenesis intermediates (also known as the “milking” way). In addition, we also noted the abundant expression of phosphate and carboxylate transporters (including the phosphate-binding protein PstS, maltose 6-phosphate phosphatase, phosphate permease, sodium-dependent dicarboxylate transporter SdcS, and bicarbonate transporter BicA) in endosymbionts (Supplementary Table 4), which might be responsible for the transportation of these sugar phosphates from symbionts.

When examining the TEM images of bacteriocyte, it is clearly showed that endosymbionts were contained in separate vacuoles designated as symbiosomes. While the presence of symbiosomes creates a suitable micro-niche for endosymbionts, it also limits their access to environmental nutrients. While we have observed high expression of ammonia consumption-related genes (top 10%) rather than the ammonia production-related genes in endosymbionts (Fig. 3C, Supplementary Fig. S6, Supplementary Table 4), the ammonia, unlike the oxygen, carbon dioxide, and methane, could yet not pass freely through the membrane of symbiosomes, and therefore must be supplied by the host. Ammonia, however, is detrimental to the host as a by-product of protein digestion and nucleotide metabolism. Here, with scRNA-seq and ST-seq data, we noted that the mussel host was massively transcribing ammonium transporter gene *RHBG-A* in bacteriocytes under normal conditions (Fig. 3C), which might facilitate the transport of ammonia into symbiosomes. In favor of the speculation, we noted that RHBG-A proteins co-located with endosymbionts (Fig. 3C) while the endosymbionts were abundantly transcribing ammonium transporter genes (the top 10% of highly expressed genes in the InS group). While the endosymbionts were also transcribing genes involved in the glutamine synthetase/glutamate synthase cycle (Supplementary Table 4), the employment of ammonium transporter could be more direct and efficient, and was also used in cnidarian-algal symbiosis ^31,32^.

Another interesting finding is that we found the also observed that the expression level of these genes involved in host-symbiont metabolism interactions, including *CYP39A1*, *RHBG-A*, and *SLC37A2*, were markedly downregulated in bacteriocytes of DeC group when methane supply was limited and when symbiont abundance dropped (Supplementary Fig. S7). These findings highlighted the possibility that the mussel hosts were interacting with symbionts dynamically based on the environments and symbiont abundance.

### Coordinated regulatory networks guide molecular functions in bacteriocytes

While the above results collectively show the host optimizing its immune and metabolic processes to facilitate symbiosis, we questioned whether there were coordinated regulatory networks guiding these processes. It was speculated that such regulatory networks would contain the majority of aforementioned immune and metabolic genes, which could share a coordinated expression pattern across all cell types and samples. We then constructed a regulatory network using weighted gene co-expression network analysis (WGCNA) of scRNA-seq data obtained on all the cell clusters in fully-symbiotic (InS group) and decolonized mussels (DeC group). Among all co-expression modules, the Mod12 modules were of interest because they contained 112/199 marker genes of bacteriocytes (Supplementary Fig. S8A, Supplementary Table 5, Supplementary Table 6). Function enrichment analysis further showed that these 112 element marker genes were involved in the immune and metabolic processes of bacteriocytes as well as cell differentiation and development (Supplementary Fig. S8B, Supplementary Table 7), certificating that Mod12 could be the core regulatory networks in the function and development of bacteriocytes.

We then screened the highly expressed transcription factors and signal transducers in the regulatory network, which may function as hub regulators of bacteriocyte function. Within the regulatory networks of bacteriocytes, we identified seven transcription factors (zinc finger protein *ZNF271*, ETS-related transcription factor *ELF-3*, autism susceptibility gene *AUST2*, histone-lysine N-methyltransferase *EZH2*, GATA zinc finger domain-containing protein *GATAD14*, hepatocyte growth factor receptor *MET*, and myoneurin) (Fig. 4A). These seven transcription factors were abundantly expressed in bacteriocytes as marker genes and correlated with the expression of 107/199 marker genes (including the aforementioned metabolic genes such as 24-hydroxycholesterol 7 alpha-hydroxylase *CYP39A1*, sugar phosphate exchanger *SLC37A2*, and ammonium transporter *RHBG-A*) and immune genes (such as *TLR2*, rabenosyn-5, lysosomal-trafficking regulator, and cathepsins) (Fig. 4A). Among these seven transcription factors, four genes (*ZNF271*, *ELF-3*, *AUST2*, and *EZH2*) were also vigorously downregulated during decolonization in conjunction with the decreased expression of another 63 marker genes (Fig. 4A). These findings highlight the regulatory role of marker gene expression in bacteriocytes. Moreover, strong interconnectivity (weight higher than 0.3) was observed between these transcription factors, certificating the coordination within these hub transcription factors when guiding the function of bacteriocytes.

**Fig. 4.**
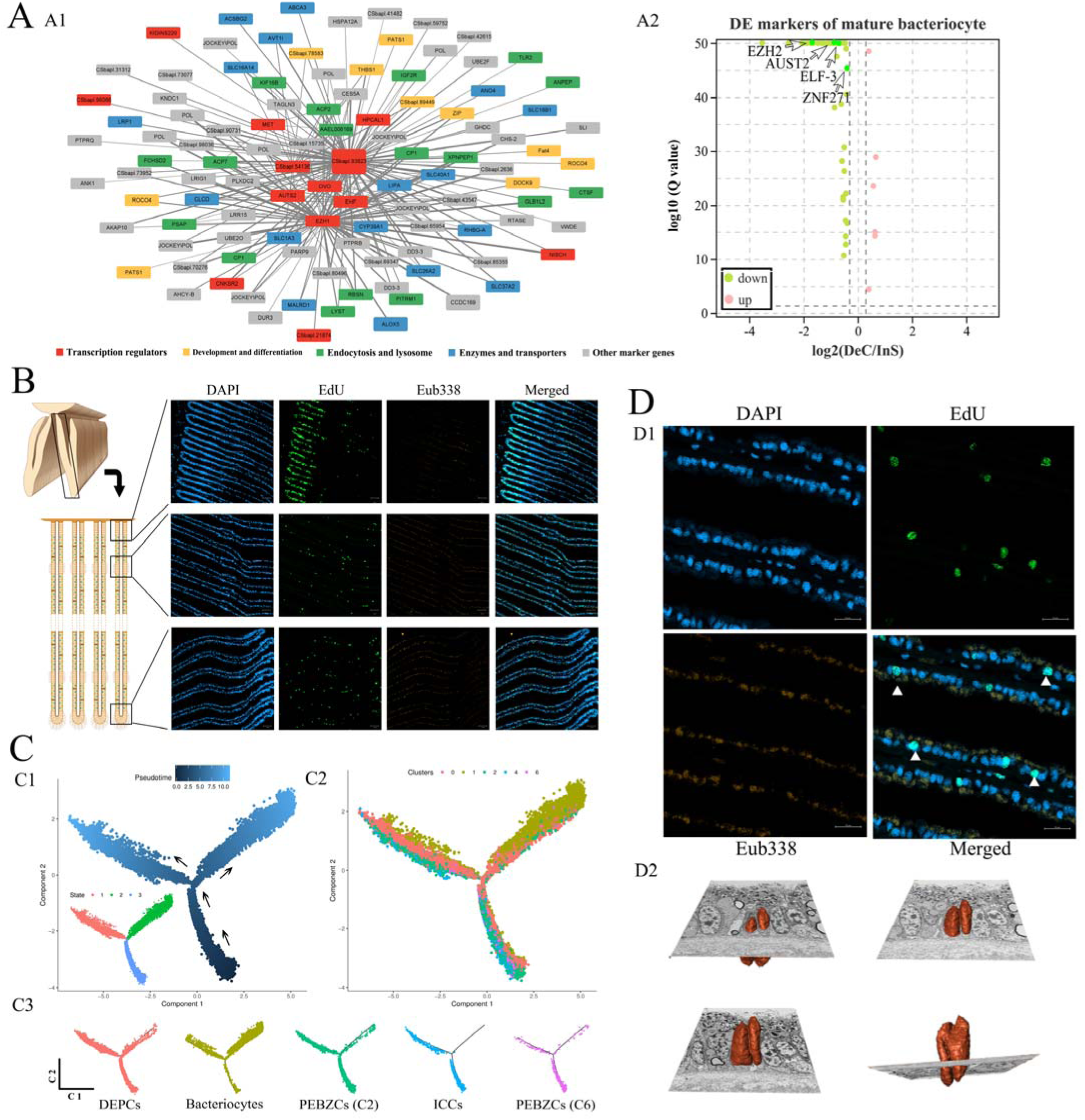
Development trajectory of deep-sea mussel bacteriocyte lineages. (A) Genetic regulatory network of bacteriocytes constructed by weighted gene co-expression network analysis (WGCNA) using scRNA-seq data. Marker genes with similar function or in same biological process are labeled in the same color (A1). Several hub transcription factors of the genetic regulatory network were found downregulated in the DeC group, accompanied by the differential expression of other 63 element markers (A2). (B) Distribution pattern of stem cells in the InS group as revealed by a 5-ethynyl 2’-deoxyuridine (EdU) staining assay. Newly synthesized DNA is labeled with EdU signals (green) while cell nuclei are stained by DAPI (blue) and symbionts are labeled using a Cy3-labeled Eub338 probe (orange). (C) Development trajectory of bacteriocyte lineages. Pseudo-time analysis in Monocle shows the successive development process from stem cells to bacteriocytes (C1), which could be further divided into three distinct states (C2). Stem cells and ciliated cells are distributed mostly in states 3 and 1 (C3) while bacteriocytes are mostly distributed in state 2. (D) Cell proliferation of bacteriocytes observed by EdU assay and 3D electron microscopy. EdU signals indicating DNA replication are observed in the nucleus region of some bacteriocytes of the InS group (white triangle, D1). Nuclear division in bacteriocytes could also be observed with 3D reconstruction of gill tissue (a total of 91 serial images with thickness at 100 nm per section).

### Successive trajectory of the development of bacteriocyte lineages

Without culturable samples of post-larval and juvenile deep-sea mussels, mechanism beneath the development process of bacteriocyte lineages remains largely elusive. The identification of marker genes of bacteriocytes and stem cells, however, provide a unique opportunity to reveal this issue. As reported in our previous study, there are three types of stem cells in gills of *G. platifrons*, including PEBZCs, DEPCs and VEPCs. By conducting EdU-labeling assay, we confirmed the proliferating activity of DEPCs, which were located at gill base (specifically at the dorsal part of the gill connected to the mantle, with positive EdU signals). Meanwhile, we also noticed that some gill cells dispersed along the gill filaments retained proliferating activity (Fig. 4B) in a manner similar to that observed in fish gills ^33^. In addition to the previously identified markers gene, our scRNA-seq and ST-seq data further identified massive cell cycle-, proliferation-, and differentiation-related genes in PEBZCs (cluster 2/6) and DEPCs (cluster 0) [including several markers of stem cells such as DEK, Serine/arginine-rich splicing factor 1, protein hedgehog, nesprin-1, protein crumbs-like 2, repulsive guidance molecule A, G1/S-specific cyclin-E gene, calcium/calmodulin-dependent protein kinase type II, prospero homeobox protein 1, Neuroglian, SWI/SNF-related matrix-associated actin-dependent regulator of chromatin subfamily D member 1, bone morphogenic protein type 2 receptor, and protein strawberry notch homolog 1] (Supplementary Table 2). Using an expression atlas of stem cells and bacteriocyte lineages, we then characterized the bacteriocyte development trajectory using three different models: pseudo-time analysis by Monocle (Fig. 4C) and partition-based graph abstraction (PAGA) analysis by Scanpy and RNA velocity analysis (Supplementary Fig. S8C, D). Results by Monocle show that the gill cells were distributed at three distinct pseudo-time states in which most stem cells, as the starting point of differentiation, were in state 3 and 1 of the trajectory (92.02%, 3.29% and 4.69% of cluster 6 PEBZCs were in state 3, 2, 1 respectively; 71.05%, 3.73% and 25.22% of cluster 2 PEBZCs were in state 3, 2, 1 respectively; 28.44%, 7.04% and 64.52% of cluster 0 DEPCs were in state 3, 2, 1 respectively). Conversely, as differentiation endpoints, approximately 88.19% of bacteriocytes were in state 2 while only 2.52% were in state 3, and 9.29% were in state 1 (Fig. 4C). In addition, as comparison, the ciliated cells (e.g, cluster 4 ICCs) were exclusively in state 3 (59.78%) and state 1 (40.13%). A similar result was also observed in PAGA analysis, in which PEBZCs were found in a more primitive state than DEPCs, ICCs and bacteriocytes (Supplementary Fig. S8C). These distinct pseudo-time trajectory states collectively confirmed developmental heterogeneity within gill cells and suggested the bacteriocytes as one of the most differentiated cells in gill. Besides, it is noticeable that a succession of processes occurred in differentiation (cell type transitions from stem cells to bacteriocytes) and maturation (subtype transitions from state 3 to state 1, or state 1 to state 2) of bacteriocytes (Fig. 4C, Supplementary Fig. S8C).

To further identify the differentiation processes in bacteriocyte lineages, we then examined the abundantly expressed genes in different pseudo-time states (Supplementary Table 8). We speculated that progenitor states would encode several hub marker genes (especially transcription factors) of descendent cells, while the expression levels of marker genes would increase during further maturation, showing maximal expression levels in the function-matured state. Our results indicate that bacteriocytes in state 2 were in the function-matured state, transcribing more marker genes (including hub marker genes) compared with the levels transcribed in the rest of the states (Supplementary Fig. S8D, Supplementary Table 8). Bacteriocytes in state 1 were in function-maturing state and showed moderate expression of marker genes. Interestingly, we also noted that most of the genes abundantly expressed in state 2 PEBZCs (cluster 2: 100/314, cluster 6: 29/32) and DEPCs (129/755) were the marker genes expressed by bacteriocytes (a total of 199 markers); also, 29/43 of DEPCs marker genes were abundantly expressed in state 1 PEBZCs (cluster 2) (Supplementary Fig. S8DE, Supplementary Table 8). These findings suggest that the state 2 cells of both PEBZCs and DPECs may have been progenitors of bacteriocytes, while DEPCs may have descended from state 1 PEBZCs. The conclusion was also supported by RNA velocity analysis, in which a positive velocity from PEBZCs and DPECs to bacteriocytes was observed in corresponding cells of adjacent region (Supplementary Fig. S8C). We also observed that state 3 bacteriocytes encoded multiple DNA replication- and cell cycle-related genes (Supplementary Table 8), indicating that bacteriocytes could proliferate directly. In support of this hypothesis, we observed increased proliferation-related signaling in several bacteriocytes, as assessed using our EdU-labeling assay and 3D electron microscopy (Fig. 4D, Supplementary Video.2).

### Co-option of conserved transcription factors in bacteriocyte development

While the development of symbiotic cells and organs may be a highly organized process, the regulatory networks guiding these developmental processes are difficult to identify in either model and non-model symbiotic associations. In this study, using our bacteriocyte developmental trajectory, we explored the mechanisms underlying bacteriocyte differentiation and maturation while focusing on the genes abundantly expressed in the progenitor state. By conducting t-Distributed Stochastic Neighbor Embedding (t-SNE) analysis, we noted a co-overlap in the t-SNE distribution of state 1 bacteriocytes and state 2 cells of DEPCs/PEBZCs and a co-overlap between state 1 cells of DEPCs and state 1 cells of PEBZCs (Fig. 5A), reconfirming that the state 2 cells of DEPCs/PEBZCs progenitor were the progenitor cells of bacteriocytes. Specifically, our results indicate that mRNA transcripts of histone-lysine N-methyltransferase *EZH2*, one of the hub marker genes in the regulatory network of bacteriocytes, were abundantly expressed in state 2 of all types stem cells (Fig. 5B, Supplementary Fig. S8F, Supplementary Table 8). Additionally, the expression of dedicator of cytokinesis protein 11, acid phosphatase type 7, cathepsin D, cathepsin L and prosaposin, which serve as element marker genes in the bacteriocyte regulatory network, and as crucial regulators of cell differentiation and lysosomal function, was also highly upregulated in state 2 of all types stem cells (Supplementary Fig. S8F, Supplementary Table 8). These genes may collectively promote the differentiation of state 2 stem cells into bacteriocytes. In addition, the expression of another three hub transcription factors in the regulatory network of bacteriocytes (*ZNF271*, *ELF-3*, and *MET*), was also robustly promoted in state 2 DEPCs (Supplementary Fig. S8F, Supplementary Table 8), thereby contributing to the differentiation of DEPCs into bacteriocytes. Similarly, we observed a robust expression of *myoneurin*, *ZNF271*, *ELF-3 and MET* in state 2 of cluster 2 PEBZCs (Supplementary Fig. S8F, Supplementary Table 8). These genes may facilitate the transition of stem cells into bacteriocytes by transducing intracellular signals and activating the expression of bacteriocyte-specific genes.

**Fig. 5.**
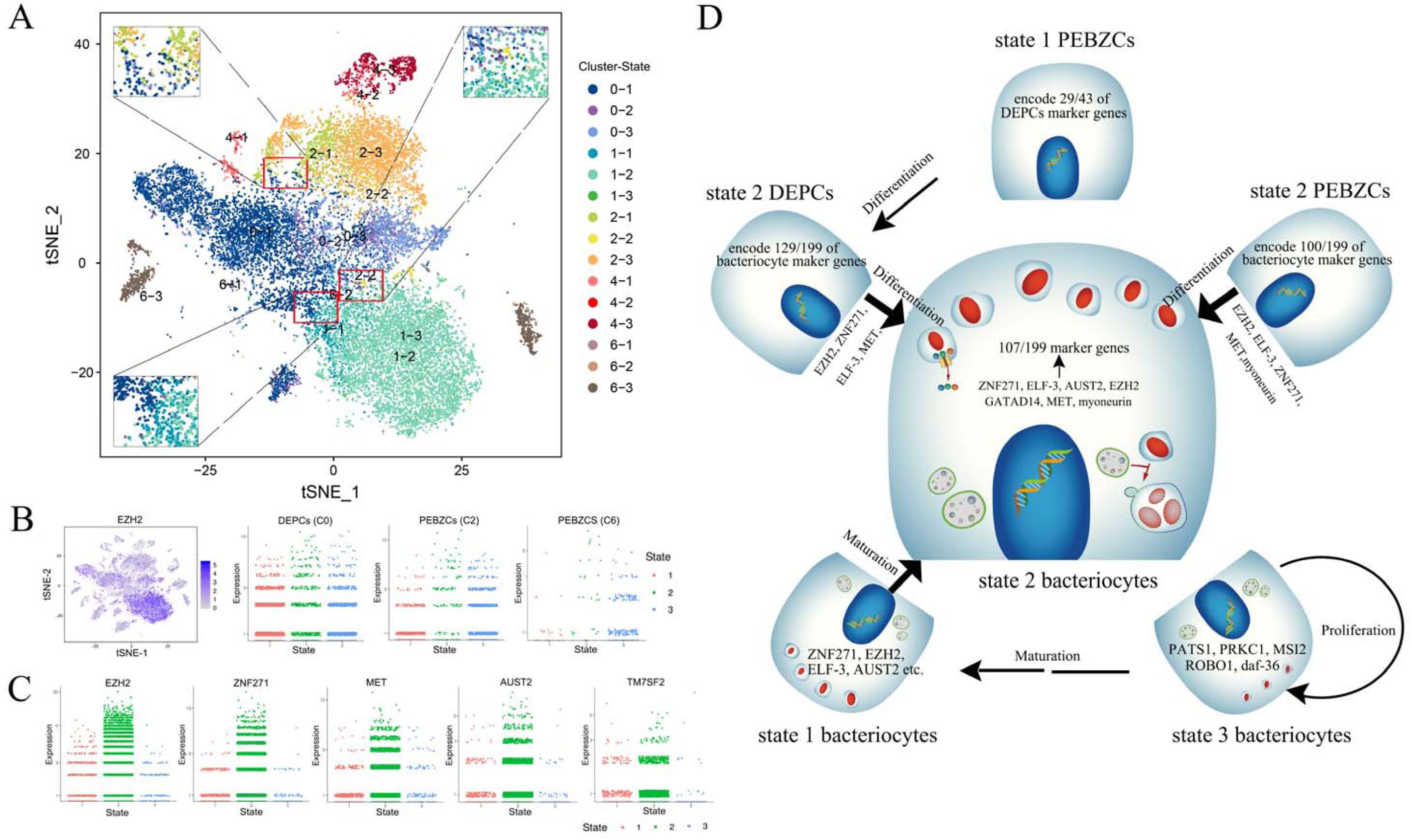
Co-option of conserved transcription factors in function and development of bacteriocytes. (A) A t-SNE plot of stem cells and bacteriocytes that in different pseudo-time states. Co-overlaps in the t-SNE distribution of state 1 bacteriocytes and state 2 cells of DEPCs/PEBZCs and between state 1 cells of DEPCs and state 1 cells of PEBZCs were observed (shown in magnified fields). (B) Expression pattern of the hub transcription factors EZH2 in color-coded t-SNE plots (left) and different states of stem cells. (C) Expression pattern of hub transcription factors in different states of bacteriocytes. (D) Schematic diagram shows function and development of bacteriocyte lineages guided by co-option of conserved transcription factors identified from the genetic regulatory networks. Arrows represent the deduced development trajectory (including cell type transition and maturation) in bacteriocytes.

During further maturation of bacteriocytes, we observed a gradual increase in the expression of four hub marker genes (*ZNF271*, *EZH2*, *MET,* and *AUST2*) from state 1 to state 2 (Fig. 5C, Supplementary Table 8). Additionally, 21 transcription factors (or activators/repressors) were also highly expressed in state 1 bacteriocytes (Supplementary Fig. S8F, Supplementary Table 8). These genes may collaboratively promote function maturation in bacteriocytes. Interestingly, we also observed that delta(14)-sterol reductase (*TM7SF2*, EC:1.3.1.70) was abundantly expressed in state 1 (Fig. 5C, Supplementary Table 8). The *TM7SF2* gene plays crucial role in the sterol metabolism of mussel host by transforming symbiont-deprived 4,4-dimethyl-cholesta-8,14,24-trienol into 14-Demethyl-lanosterol, implying the contribution of symbiont-deprived nutrients in bacteriocyte maturation.

While the aforementioned hub transcription factors and signaling transducers may be the pioneer molecules promoting bacteriocyte development, we questioned whether they were evolutionarily novel genes diverged from non-symbiotic ancestors or conserved across evolution. Our phylogenetic analysis of protein sequences obtained from homologues of these genes in Lophotrochozoa indicated that all of these genes were highly conserved across the mollusk and in harmony with the general host phylogenies (Supplementary Fig. S9). We therefore conclude that bacteriocyte development occurred via co-option of conserved genes rather than via evolutionarily novel genes (Fig. 5E).

## Discussion

In addition to being an evolutionary novelty, symbiotic cells and organs provide a suitable niche for symbiosis and endow holobionts with a unique adaptation to surrounding environments ^6^. In this study, we integrated the data obtained on single-cell omics, phagocytosis assay, EdU labeling assay and 3D electron microscopy to uncover the function and development of bacteriocytes in deep-sea mussels. Our findings indicate that these processes were guided by an ancestral intrinsic toolkit via co-option of conserved transcription factors and modulation by endosymbionts. Our results show how both partners interact with each other closely and shape the function and development of symbiotic cells cooperatively, which is crucial in understanding the evolution of chemosymbiosis and the adaptation of holobionts in habitats such as the deep sea ^34,35^.

The establishment and maintenance of endosymbiosis with exogenous bacteria is challenging for multicellular organisms because of the host immune system, which requires an adjustment that entails an inoculation with, and proliferation of, symbionts inside the cell ^36^. Using a single-cell expression atlas, we characterized the immune strategy adopted by bacteriocyte lineages, and observed that only a few PRRs were abundantly expressed in bacteriocytes despite their expansion in the deep-sea mussel genome ^18^. While these PRRs might participate in other process, such as early symbiotic acquisition in juveniles and immune defense again pathogenic bacteria, our findings suggested that only a few PRRs are needed to maintain symbiosis in normal bacteriocytes of adult mussels. Nevertheless, the bacteriocytes are still cable to phagocytize exogenous bacteria even when they were fully colonized by endosymbionts, as observed by our phagocytosis assay and some recent studies ^37^. The phagocytotic ability of gill cells may be conserved in most mollusks, including non-symbiotic ones ^37–39^, which directly facilitate the entrance of endosymbionts in deep-sea mussels and may contribute importantly to the establishment of endosymbiosis in other mollusks. The phagocytosis of exogenous bacteria, however, also raise questions on how mussels discriminate non-symbionts from symbionts, and symbionts in bad states from good states inside the phagosomes or symbiosomes. It is often observed that the bacteriocytes digest endosymbionts via lysosome. Although the lysosome-mediated digestion could provide nutrients to hosts (“farming” way), we rarely observed proliferating endosymbionts or free symbionts that being engulfed in bacteriocytes of adult mussels, as evidenced by EdU labeling assay and 3D electron microscopy. The contradicts in the number of new born endosymbionts and digested endosymbionts suggested that lysosome-mediated digestion could play other roles in addition to providing nutrients. Recently, it is showed that the digestion of symbionts could be regulated by mTORC1 through nutrient signaling pathway and promoted under reduced nutrient supply (either by death of endosymbionts or removal of methane) ^40–42^. We therefore speculated that lysosome-mediated symbiont digestion may be a secondary option to obtain nutrition under normal circumstance, but an efficient way to control the symbiont population and obtain nutrition under abrupt stresses or other emergencies. In support of this speculation, we observed a significant repression of lysosome activity in mussels after long-term methane starvation. These findings further certificate that symbiotic associations in deep-sea mollusks are nutrition driven and imply the metabolic interaction between mussel hosts and symbionts ^11^.

Since their first characterization, how deep-sea mussels acquire nutrients from symbionts have been hotly debated. Besides the digestion of symbionts, it is also suggested that the mussel hosts could acquire nutrients via the receipt of secreted metabolites from symbionts (“milking” way) ^37^. However, the exact metabolites and how they are transported between symbionts and bacteriocytes, have remains unclear and difficult to characterize ^18,19,22,30,43^. Genomic information showed that the methanotrophic endosymbionts could provide sterol intermediates to the mussel hosts. Recently, Geier et. al have also identified several specialized metabolites from the host-microbe interface with metaFISH and AP-MALDI-MSI analysis ^22^, which are specific to the mussel-endosymbiont interaction. With the help of state-of-the-art single-cell sequencing, we here showed that deep-sea mussels have remarkably reshaped bacteriocyte metabolism to maximize symbiotic profits for both partners. For example, bacteriocytes encode dozens of genes involved in the biosynthesis and transport of carbohydrates, lipids, amino acids, and vitamins to improve the acquisition of nutrition from symbionts. Noticeably, we have demonstrated that the bacteriocytes are massively transcribing a sugar phosphate exchanger gene, which co-locates with symbionts and might help to retrieve fructose-6P directly from methanotrophic symbionts. Fructose-6P is believed to be converted into glucose-6P, fructose-1,6P_2_, and glyceraldehyde-3P, and further supports the gluconeogenesis and TCA cycles of both mussel hosts and symbionts. In addition, we also showed that the mussel hosts could supplies ammonia to symbionts directly via ammonium transporters, which greatly increases the efficiency and profits of symbiosis and is rarely reported ^32^. Another interesting but rationale finding is that bacteriocytes are abundantly expressing enzymes involved in the turnover of sterol intermediates, as the symbionts are the main source of these intermediates. While there might be more metabolites that are supplied by, or transported between, symbionts and bacteriocytes, our findings demonstrate that the scRNA-seq could serve as a suitable tool to address such issue in non-model holobionts.

As a specialized and stable niche for endosymbiosis, where and how bacteriocytes come from are intriguing and crucial in understanding the evolution of symbiosis ^2,6^. While previous studies demonstrated that the stem cells in the growth zones of gill could differentiate into bacteriocytes, our single-cell transcriptome data has for the first time traced the successive development trajectory of bacteriocytes with molecular evidences. We showed that the stem cells and bacteriocytes could be further divided into different molecular states and only a small subset of stem cells are the progenitors of bacteriocytes. Besides, the differentiation and maturation of bacteriocytes could be guided by a set of mollusk-conserved transcription factors (including *EZH2*, *ELF-3*, *ZNF271*, and *AUST2*), following the gradual increase of these genes from stem cells to bacteriocytes. The synergistic modulation shown by these transcription factor genes confirmed that the formation of symbiotic cells in deep-sea mussels occurred via co-option of conserved, rather than evolutionarily novel genes ^44^. Interestingly, similar phenomena are also observed in the deep-sea scaly-foot snail, in which the formation of biomineralized skeleton is driven by an ancestral intrinsic toolkit that is conserved across the mollusk ^45^. These findings collectively demonstrate the developmental plasticity of the mollusk. Noticeably, we also demonstrated that the key regulatory networks responsible for the development of bacteriocytes were also controlling the immune and metabolic process, highlighting the cooperation in the function and development of bacteriocytes. An intriguing finding is some bacteriocytes away from the growth zone are proliferating, as evidenced by both EdU assay and 3D electron microscopy analysis. It yet remains to be elucidated whether these cells are new born bacteriocytes from stem cells dispersed along gill filaments or are able to divide after “reprograming”. In favor of the second speculation, we noted that about 2.52% of bacteriocytes are abundantly expressing DNA replication- and cell cycle-related genes. Another interesting finding is that the function and development of mussel bacteriocytes may have been influenced by symbionts via sterol metabolism. Symbiont participation in the development of symbiotic cells and organs has attracted increasing attention for several years, but has remained largely unclear at the mechanistic level ^46,47^. Sterol metabolism in symbionts may be a common mechanism in the development of symbiotic cells and organs since several animal hosts rely on their symbionts for sterol intermediates ^48–50^. In support of this theory, we noted a suppression of sterol/steroid biosynthesis in the symbiont-depleted deep-sea mussels after long-term atmospheric cultivation without methane supply ^42^; furthermore, a crucial sterol metabolic gene was differentially expressed during the maturation of bacteriocytes, highlighting possible participation of sterol metabolism in the phenotypic plasticity of gill tissue ^42^. Other studies have also shown the modulation of host development and reproduction by symbiotic *Wolbachia* via steroid-nuclear receptor signaling pathway ^51^. Besides the sterol metabolism, it is also noteworthy that the glucose and ammonia metabolism could also serve as intracellular signals of symbiosis-related process, such as mTOCR1-mediated symbiont digestion, and controls the population of symbionts ^52–55^. While the regulatory role of symbionts in the function and development of symbiotic cells remains to be fully elucidated, our findings highlight the robust plasticity of molluscan bacteriocytes, which assist in the distribution of mollusk across a wide range of habitats including the deep sea.

There are also some limitations for the present study. For example, some hypotheses raised by the present study still needs to be verified in future studies, using mussel samples in different symbiotic states (including aposymbiotic and recolonized mussels as well as post-larval and juvenile mussels). Nevertheless, by combing high-resolution single-cell transcriptome data, phagocytosis assay, EdU assay and 3D electron microscopy data, we have successfully set up pipeline to reveal the molecular function and development process of bacteriocyte lineages in non-model deep-sea mussels. Our results show that the function and development of bacteriocytes were highly coordinated via co-option of conserved genes and could be dynamically modulated by symbionts. The intimate symbiotic associations and robust plasticity of symbiotic cells have rendered the deep-sea mussel one of the most successful organisms in the deep sea.

## Materials and Methods

### Experimental design

The goal of the present study was to address the molecular function and developmental trajectory of bacteriocytes in the deep-sea mussel *G. platifrons*. To achieve this goal, we employed methanotrophic *G. platifrons* collected from cold seeps as our model, and used the spatial transcriptome, single-cell transcriptome, and meta-transcriptome to characterize both the bacteriocytes and endosymbionts. We also used an *in situ* transplantation assay to construct the decolonized mussel to elucidate the dynamic response of symbiotic cells in response to symbiont depletion. We employed EdU-labeling, phagocytosis, *in situ* hybridization (ISH) assays, immunofluorescence (IF) assays, transmission electron microscopy (TEM), 3D electron microscopy and phylogenetic analysis to verify hypotheses obtained from the sequencing data.

### In situ transplantation and animal collection

*G. platifrons* mussels were collected from the cold seeps (22°06′N, 119°17′E) of South China Sea during the cruises of 2018 and 2020. The mussel fauna live at 1,120 m beneath the surface at a temperature of approximately 3.35°C, salinity of approximately 35.54 psu, dissolved oxygen of 2.98–3.17 mg/L, and methane concentrations of up to 31,227 ppm (seepage region) ^56^. To shelter the specimens from temperature and pressure fluctuations during sampling, all mussel samples were collected using a self-designed isothermal isobaric sampler and a self-designed manually controlled macrofauna *in situ* sampling device as described previously ^41^. In brief, a total of 20 mussels living in close proximity to active fluid seepages were collected as representatives of normal mussels (designated as the InS group). Among them, seven mussels were collected using an isothermal isobaric sampler (in 2018), seven mussels were collected using a multipurpose *in situ* sampling device (in 2020) after treating with RNAsafer stabilizer reagent (Omega Bio-Tek, Norcross GA, USA) *in situ*, and the remaining six mussels were collected using a multipurpose *in situ* sampling device after being treated with 4% paraformaldehyde *in situ*. Only one mussel from the InS group was used for single cell transcriptome sequencing (scRNA-seq, Chromium platform of 10x Genomics, Pleasanton CA, USA) and spatial transcriptome sequencing (ST-seq, Visium platform of 10x Genomics), and all 14 mussels collected by isothermal isobaric sampler or after RNA stabilizing treatment were subjected to meta-transcriptome sequencing. Mussels collected after paraformaldehyde treatment were subjected to ISH, IF, and TEM imaging. During the 2018 cruise, we also performed an *in situ* transplantation assay, in which dozens of mussels in the seepage region were translocated to an authigenic carbonate region having low concentration of CH_4_ (at approximately 100 m away from seepage, methane concentration had decreased to 800 ppm). Five of the transplanted mussels (designated as the DeC group) were retrieved 604 days later during the 2020 cruise using an isothermal isobaric sampler (one mussel, used for scRNA-seq, ST-seq and meta-transcriptome) or multipurpose *in situ* sampling device (four mussels, used for meta-transcriptome) after treatment with RNAsafer stabilizer reagent. After the retrieval and depressurization of the isothermal isobaric sampler, the mussels were instantly dissected into two halves to remove excess seawater and quickly frozen in an isopentane bath (Macklin, Shanghai, China) with liquid nitrogen for scRNA-seq and ST-seq.

### ScRNA-seq

Two individual mussels having similar size (approximately 80 mm in length) from the InS and DeC groups were used for scRNA-seq. Because of the difficulties involved in isolation and *in vitro* culture of gill cells, and because mussel sampling and single cell preparation can potentially influence gene expression, scRNA-seq analysis of *G. platifrons* gill tissue was performed using nuclei isolated from fresh frozen samples collected in an isothermal isobaric manner. The isolated nuclei were assayed per 10×Genomics single cell protocol by generating a Single-cell Gel Bead-In-EMulsion (GEM) on a GemCode single-cell instrument (https://www.10xgenomics.com/support). Full-length cDNA was synthesized using Chromium Next GEM Single Cell 3’ Reagent Kit v3.1 and sequenced on an Illumina HiSeq X Ten platform (Gene Denovo Biotechnology Co., Guangzhou, China).

### Cryosectioning and ST-seq

To help determine the taxonomy of all gill cells, especially bacteriocytes, we also conducted ST-seq with the cross-sectioned middle region of the gill (ventral view) from the InS group. The gill tissue used for 10×Visium ST-seq was first incubated with precooled methanol (4°C) for 15 min to reduce potential damage by the cryostat blade during cryosectioning. The tissue was then embedded in OCT (Sakura, Torrance, CA, USA) and cross-sectioned at the thickness of 10 µm through the middle part of the gill using a cryostat (Leica CM1950, Heidelberger, Germany). Gill sections were then stained with 4,6-diamidino-2-phenylindole (DAPI, Thermo Fisher, Waltham, MA, USA) and HE (Sangon Biotech, Shanghai, China), and morphology of the gill-tissue sections was assessed under light microscopy (Nikon ECLIPSE Ni, Tokyo, Japan). Successive sections, obtained from the same blocks of embedded tissue, were placed on Visium Spatial Tissue Optimization Slides within the capture area for tissue optimization assay. For the sequencing assay, gill sections were first mounted on Visium Spatial Gene Expression Slides and permeabilized according to previously described parameters (permeabilization time of 9 min) after HE staining and imaging using light microscopy. After reverse transcription and library preparation, all samples were sequenced on an Illumina HiSeq X Ten platform (Gene Denovo Biotechnology Co., Guangzhou, China). The rest sections of the same tissue were further subjected to ISH assays to assist the cell type annotation.

### Cell clustering, cell type annotation, and analysis of differentially expressed genes

Single-cell clustering of *G. platifrons* gill tissue was first performed using scRNA-seq—derived data. After quality control, all raw reads were mapped onto a *G. platifrons* reference genome. The genome was first reported by Sun et al. ^18^ and updated recently with Hi-C and high-depth PacBio long-read sequencing by us (GenBank accession NO. JAOEFJ000000000, unpublished data). Unique molecular identifiers (UMIs) in each sequenced read were counted and corrected for sequencing errors. Using valid barcodes that were identified based on EmptyDrops method, the gene matrices of all the cells were then produced and imported into Seurat (version 3.1.1) for cell clustering using a graph-based clustering approach.

Single-cell clustering of *G. platifrons* gill tissue was also performed using ST-seq—derived data, which have the advantage of cell type annotation. A slide image obtained before permeabilization was first imported into Space Ranger software for fiducial and Visium barcoded spot alignment. After decoding correlations between tissue, capture spots, and barcodes, a splice-aware alignment of sequencing reads to the *G. platifrons* genome was performed using STAR in the Space Ranger software package.

An integrated analysis of data derived using scRNA-seq, ST-seq, and *in situ* hybridization assay was then used for cell type annotation in the InS group. In brief, anchor-based integration was first used to integrate ST-seq data with scRNA-seq data using the FindIntegrationAnchors command in Seurat-v3.2. All cell-type labels in scRNA-seq were then transferred to spatial data using the TransferData command. Cell type prediction scores indicating the similarity between ST-seq spots and scRNA-seq cell clusters were calculated simultaneously, and only spot-cluster pairs with highest scores were considered for further cell-type annotation. For further annotation and cell-type verification, cell types in gill tissue were inspected using HE stained images of gill cryosections and tissue plot with spots colored by clustering of ST-seq data. Several cell markers obtained using the FindAllMarkers function in Seurat were also cloned and subjected to *in situ* hybridization using gill tissues from the InS group.

To further investigate the biological function of all identified cell clusters, differentially expressed genes (DEGs) were surveyed in both scRNA-seq and ST-seq. For scRNA-seq, expression values of all identified genes in a given cluster were compared against the rest of the cell clusters using Wilcoxon rank sum test. Genes expressed mainly in the target cluster (more than 25% of cells were designated as a target cluster), and showing at least 1.28-fold upregulated expression levels and *p*-values less than 0.01, were considered as DEGs per cell cluster. For ST-seq, the Model-based Analysis of Single cell Transcriptomics (MAST) in the R package was used to determine DEGs in a single cell cluster using the same criteria as those used for scRNA-seq. All the genes in ST-seq were further analyzed for spatially-specific DEGs using markvariogram in the Seurat R package and mark-segregation hypothesis testing in the trendsceek R package. Genes with r.metric.5 parameter values of less than 0.8 and *p*-values of less than 0.01 were designated as spatially DEGs.

### Meta-transcriptome sequencing

To explore the expression atlas of symbionts in the InS and DeC groups, we performed meta-transcriptome sequencing using the 14 samples from the InS group and five samples from the DeC group as described previously. Total RNA extraction, removal of eukaryotic and prokaryotic rRNA, and synthesis of cDNA library using bulk-seq were conducted as described previously ^41^. cDNA libraries belonging to the InS and DeC groups was finally sequenced using an Illumina HiSeq 2500 platform with paired-end reads performed by Novogene (Tianjing, China). After quality control, the filtered reads were aligned against both the endosymbiont and *G. platifrons* genomes using HISAT (v2.0.4). The genome of methanotrophic symbionts was first reported by Takishita et al., (2017) and we updated it with high-depth PacBio long-read sequencing (raw data deposited in NCBI with accession NO. PRJNA891367) ^57^. The expression levels of host and symbiont genes were calculated using HTSeq (v0.6.1), while significant differences between groups were determined using DESeq2 (v1.10.1). Additionally, the top 10% of genes with most abundant mRNA transcripts in the InS group were designated as abundantly expressed genes in order to survey the biological processes that actively occurred under normal conditions *in situ*. For PCoA analysis, Bray-Curtis distance between samples was first calculated and subjected for subsequent analysis and visualization.

### GO/KEGG analysis, cell trajectory construction, and WGCNA of bacteriocytes

Gene Ontology (GO) annotation of *G. platifrons* genes was obtained using Blast2GO software (version 5.2) and employed for GO enrichment analysis using homemade scripts. DEG numbers for every GO term were first calculated and the significantly enriched GO terms were determined using a hypergeometric test (FDR less than 0.05). KEGG pathway enrichment analysis was conducted using homemade scripts with KEGG annotations obtained from the KEGG database (http://www.genome.jp/kegg/pathway.html).

Single cell trajectory analysis was conducted using Monocle (Version 2.6.4) with reversed graph embedding algorithm and confirmed by PAGA analysis using Scanpy (v1.6.0). Gene expression matrix of stem cells and bacteriocytes was used in the analysis and visualized using the orderCells function of Monocle (sigma = 0.001, lambda = NULL, param. gamma = 10, tol = 0.001). For the PAGA analysis, the connectivity of each cell cluster was calculated based on the partition-based graph abstraction algorithm. After cell embedding with ForceAtlas2, the pseudo-time value of each cell was then calculated with the DPT algorithm. Weighted gene co-expression network analysis (WGCNA) was conducted using the WGCNA package (v1.47) in R with gene expression values of power = 7 and minModuleSize = 50, and visualized using Cytoscape (v3.8.2).

### EdU-labeling, phagocytosis, ISH assays, IF assays, 3D electron microscopy, and phylogenetic analysis

The EdU-labeling assay was carried out *in situ* using mussels collected from the seepage region. In brief, mussels were incubated with 5-ethynyl 2’-deoxyuridine (EdU, final concentration of 40 µM) in a self-designed, manually controlled macrofauna *in situ* experiment device for approximately 18 h, and then retrieved using an isothermal isobaric sampler. The Click-iT Plus EdU Imaging Kit (Thermo Fisher) was then used to visualize the EdU signal.

Phagocytosis assay was conducted using primary gill cells obtained from the fresh collected mussels using previously described methods with modifications ^38,58^. Because endosymbiotic methanotrophs are nonculturable, *Vibrio alginolyticus*, an environmental bacterium isolated from the cold seep, was used in this assay and labeled with fluorescein isothiocyanate (FITC, Sigma, St. Louis, MO, USA) before use. To collect primary gill cells, gill tissue was first treated with 1% trypsin (diluted in sterilized seawater) at 4°C for 30 min, and then successively centrifuged at 300 g (4°C, 5 min) and 800 g (4°C, 5 min). Cell pellets were resuspended in modified L15 medium (Gibco, Carlsbad, CA, USA; supplemented with 0.54 g/L KCl, 0.6 g/L CaCl_2_, 1 g/L MgSO_4_, 3.9 g/L MgCl_2_, and 20.2 g/L NaCl) to a final concentration of 1×10^6^ cells mL^−1^ and incubated with the same volume of FITC-labeled *V. splendidus* (1×10^8^ cells mL^−1^) for 30 min at 4°C in the dark. Primary cells were then washed three times with modified L15 medium to remove extracellular bacteria. Cells were stained with DAPI and DiI perchlorate, and were then imaged using a laser scanning confocal microscope (Zeiss LSM710, Jena, Germany).

ISH assays were conducted as described previously ^59^. For cell-type verification, cell markers were cloned using gene-specific primers to synthesize digoxigenin-labeled ISH probes. Fluorescent ISH analysis of symbionts was performed using a Cyanine 3 (Cy3)-labeled Eub338 eubacteria probe (5’-GCTGCCTCCCGTAGGAGT-3’) or FITC-labeled pmoB (methanotroph-specific gene) probe (5’-CGAGATATTATCCTCGCCTG-3’).

For the IF assay, unique peptide fragments of the 24-hydroxycholesterol 7 alpha-hydroxylase *CYP39A1*, sugar phosphate exchanger *SLC37A2*, and ammonium transporter *RHBG-A* were first synthesized by Sangon Biotech (Shanghai, China) and employed as antigens to produce rabbit polyclonal antibodies. After antigen affinity purification and specificity verification by ELISA and western blot, the antibodies were then used in an IF assay using paraffin embedded gill tissue (5 μm) according to methods described previously ^59^ and with the assistance of ServiceBio (Wuhan, China). Specifically, anti-lipid A antibody (ab8467, Abcam, Cambridge, MA, USA) was used to indicate endosymbionts. After incubation with FITC- and Cy3-labeled secondary antibodies (Servicebio, Wuhan, China), gill sections were visualized and imaged using fluorescent microscope.

For the 3D electron microscopy assay, gill tissues collected from adult mussels (obtained in isothermal way) were fixed by paraformaldehyde-glutaraldehyde, treated with 1.0% osmium tetroxide (OsO4) and infiltrated with acrylic resin successively. The embedded tissues were then serially sectioned with a thickness of 100 nm using an ultramicrotome (EM UC7, Leica, Vienna, Austria) and mounted onto a silicon wafer before imaging. The scanning electron microscope micrographs of mounted sections were captured using Helios NanoLab 600i FIB-SEM (FEI, Hillsboro, USA). The obtained serial section images were then aligned in Amira (version 2019.3) for the 3D reconstruction of bacteriocytes and cell proliferation analysis.

For phylogenetic analysis of hub transcription factors, homologue proteins were first obtained using NCBI blastp and aligned using Seaview. Maximum likelihood phylogenetic trees of these proteins were then constructed using Mega software (v11) in Jones-Taylor-Thornton model with bootstrap of 100.

## Acknowledgments

We thank the crewmembers of the R/V Kexue for their assistance in sample collection, and all the laboratory staff for continuous technical advice and helpful discussions. We also thank Prof. Qiang Lin from the South China Sea Institute of Oceanology, Chinese Academy of Sciences for insightful comments and suggestions that improved the manuscript.

## Funding

The present study was supported by the National Natural Science Foundation of China (42106134, 42030407, and 42076091), the Key Research Program of Frontier Sciences (ZDBS-LY-DQC032) and Strategic Priority Research Program of the Chinese Academy of Sciences (XDA22050303, XDB42020401), Key Deployment Project of Centre for Ocean Mega-Research of Science, CAS (COMS2020Q02) and Marine S&T Fund of Shandong Province for Pilot National Laboratory for Marine Science and Technology (Qingdao) (No. 2022QNLM030004).

## Author contributions

Conceptualization: H.C., Chaolun Li; Sample collection: M.W., M.L., Z.Z., Chao Lian., L.Z.; Environment investigation: M.W., L.C.; Transcriptome analysis: H.C., Z.Z., M.L.; EdU labeling, phagocytosis, ISH and IF assays: H.C., M.L., G.H., Z.Z., H.W.; Phylogenetic analysis: H.C., H.Z.; Supervision: C.L.; Writing—original draft: H.C.; Writing—review & editing: H.C., M.W., Chaolun Li.

## Conflict of interest

The authors declare that they have no competing interests.

## Data Accessibility

All data needed to evaluate the conclusions in the paper are present in the paper and/or the Supplementary Materials. All sequencing data were deposited in the NCBI BioProject database (https://www.ncbi.nlm.nih.gov/bioproject/) under accession NO. PRJNA838712. Raw images for 3D electron microscopy were deposited in Figshare (https://figshare.com/articles/figure/Raw_image_for_3D_electron_microscopy_of_mussel_gill/23575734).

## Notes

### Competing Interest Statement

The authors have declared no competing interest.

### Summary of Updates

Text revision on the introduction, result and discussion part.

