## Supplementary figures and images for "Cooperation between bacteriocytes and endosymbionts drives function and development of symbiotic cells in mussel holobionts"

### Supplementary Fig. S

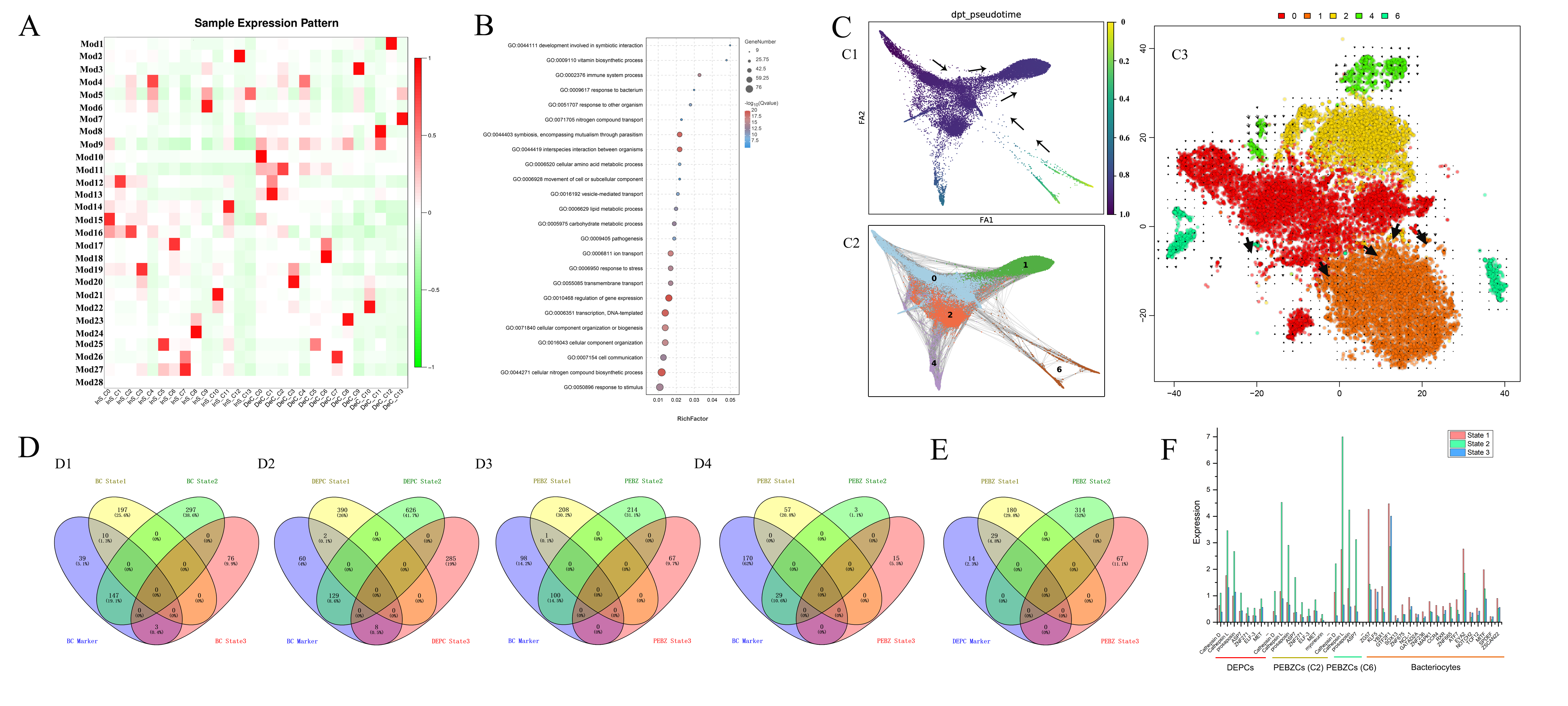

### Supplementary Fig. S1

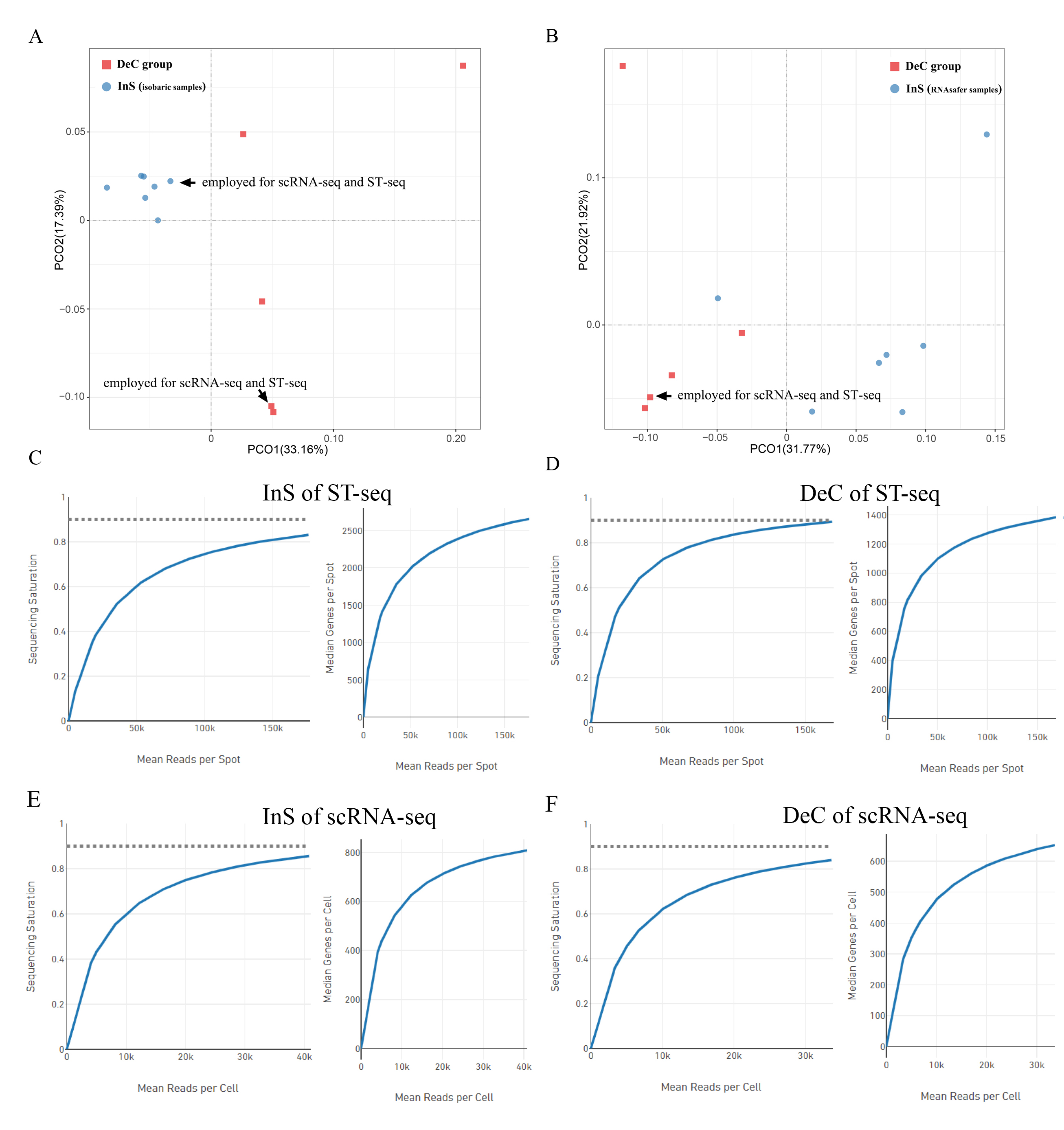

### Supplementary Fig. S2

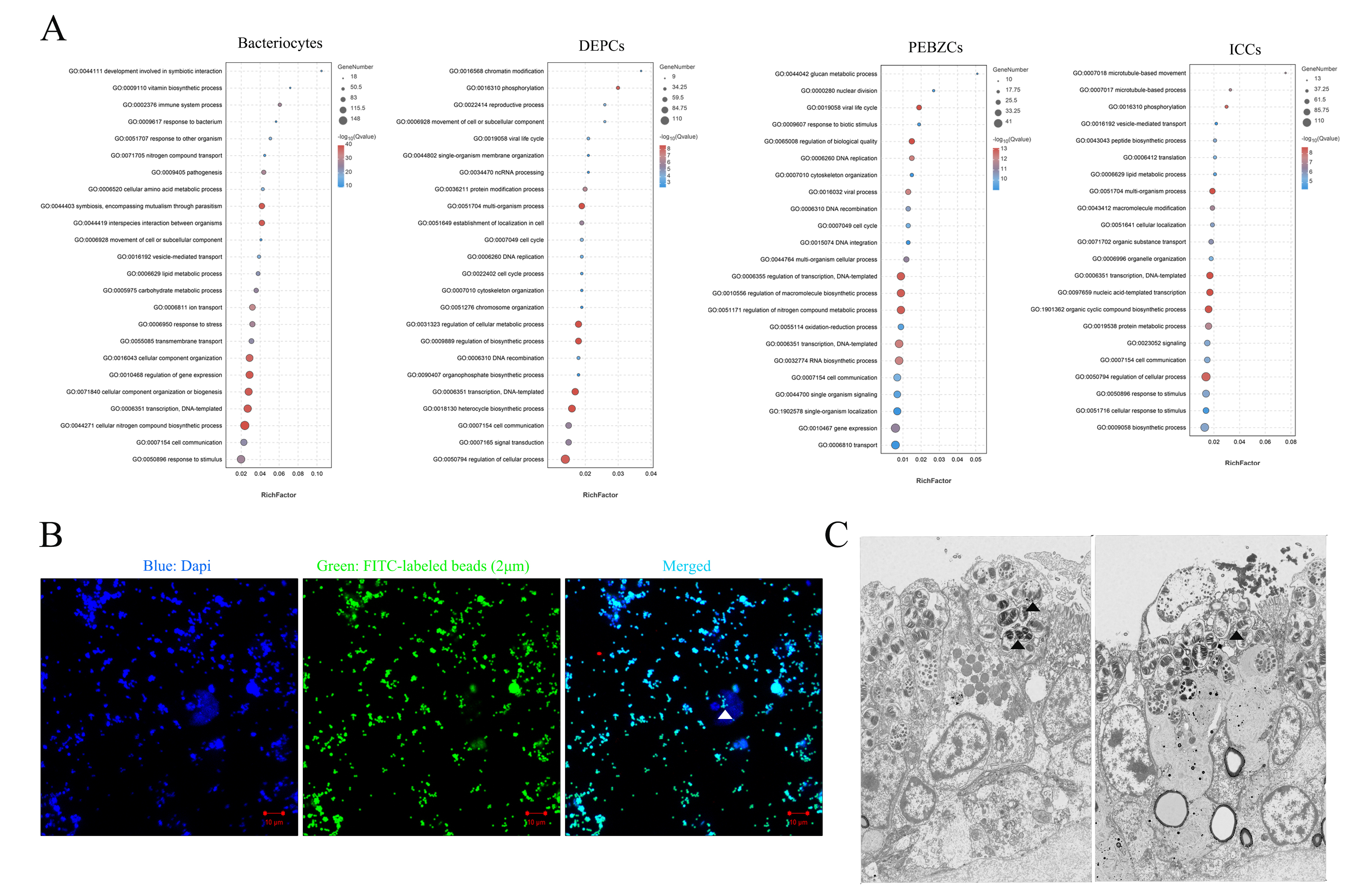

### Supplementary Fig. S3

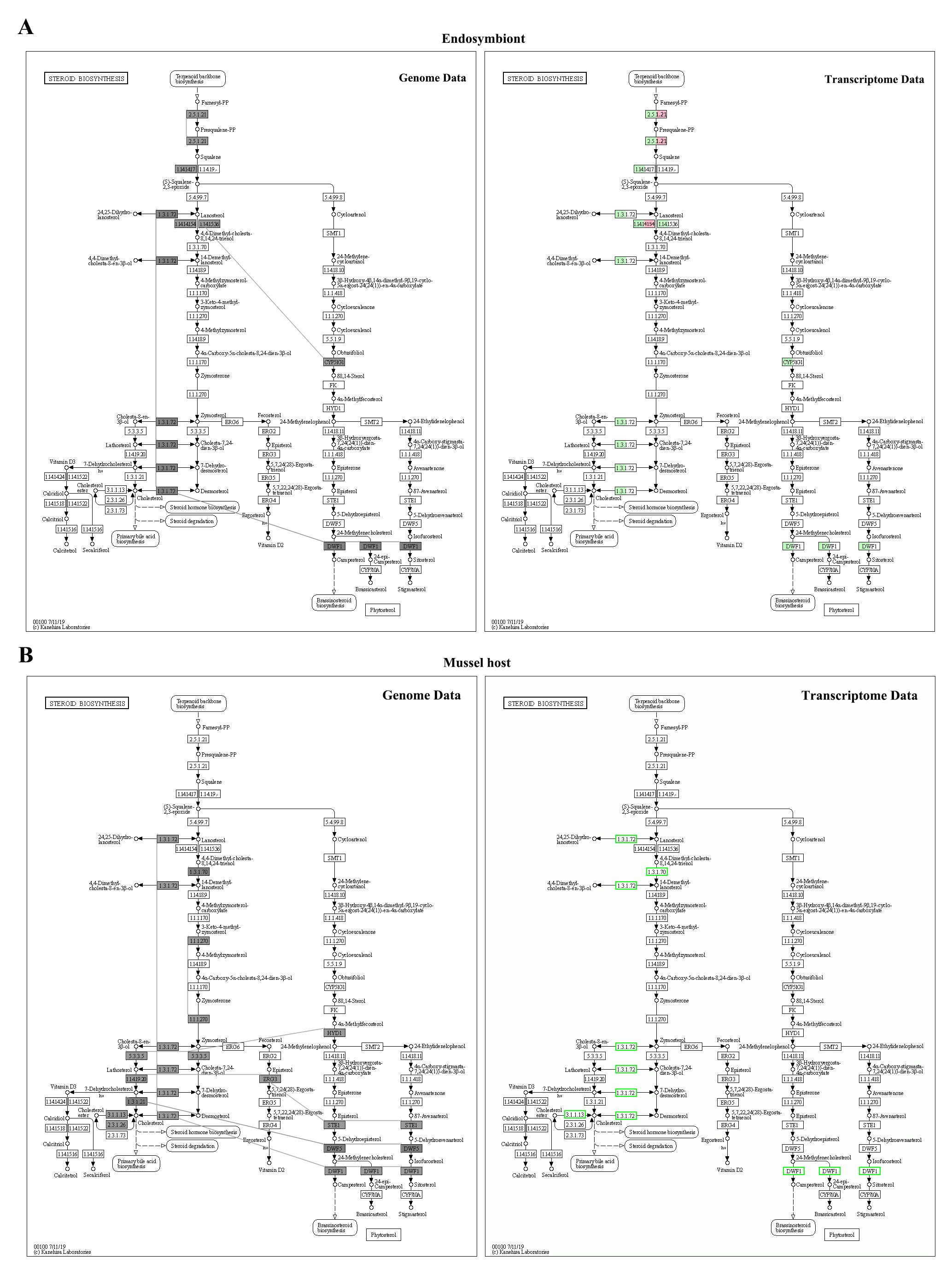

### Supplementary Fig. S4

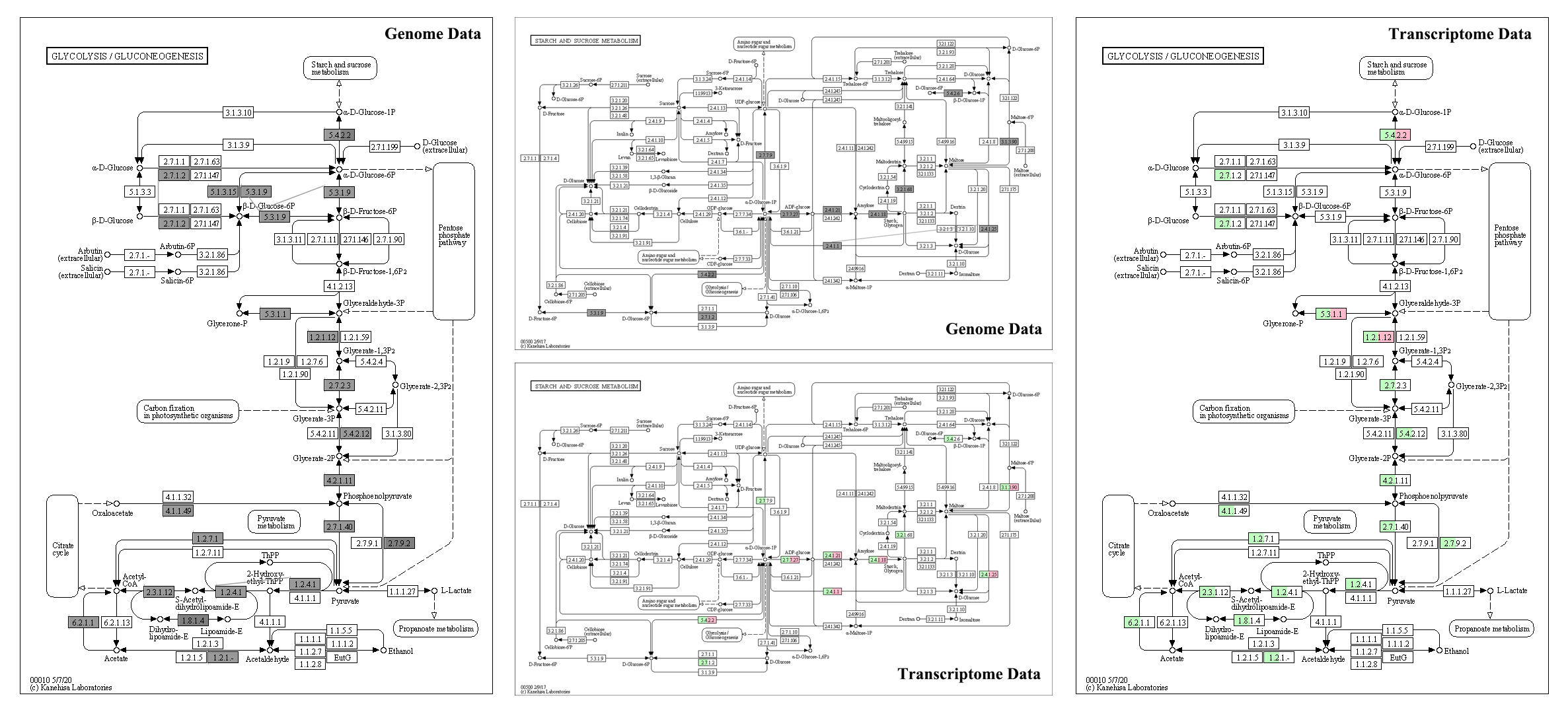

### Supplementary Fig. S5

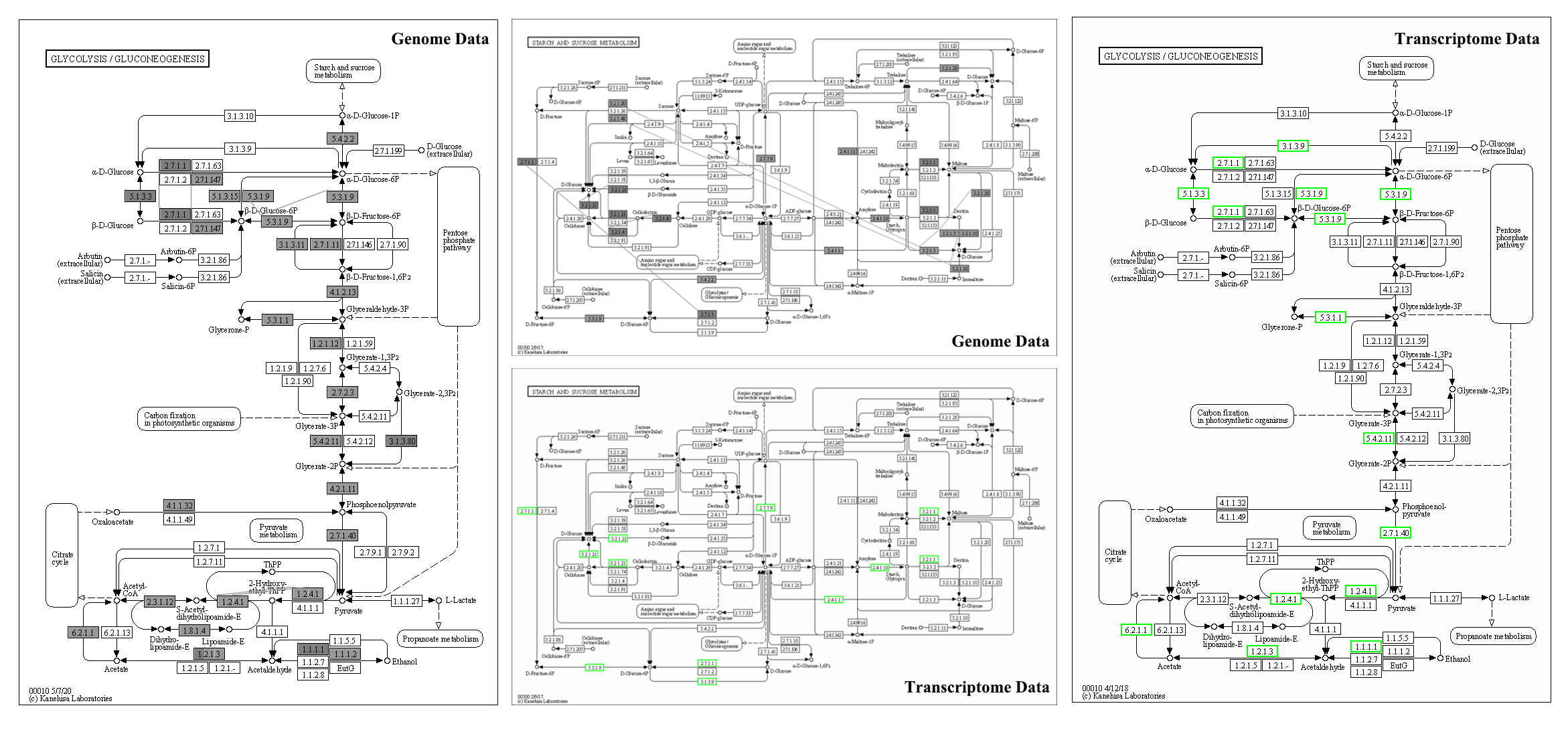

### Supplementary Fig. S6

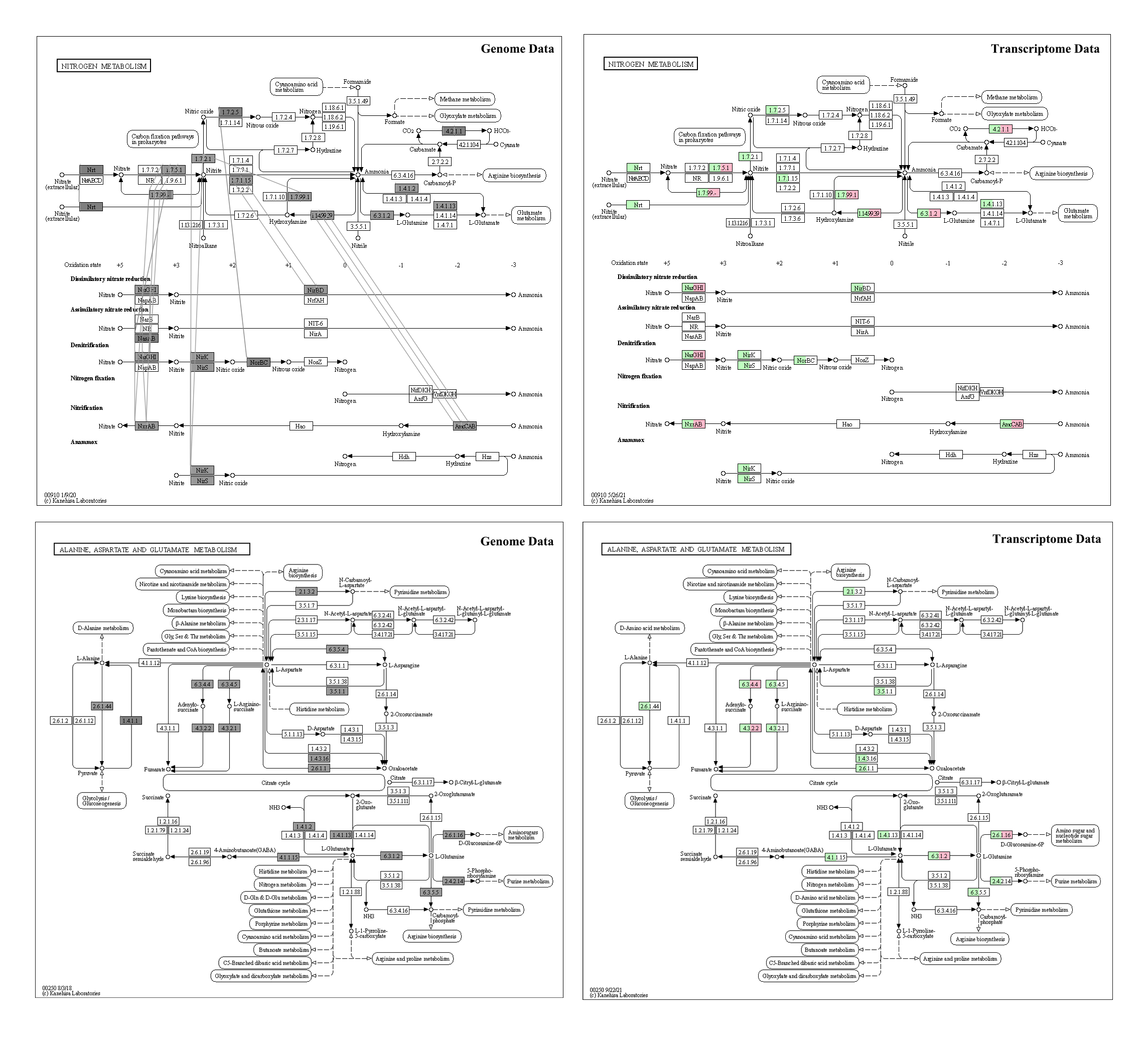

### Supplementary Fig. S7

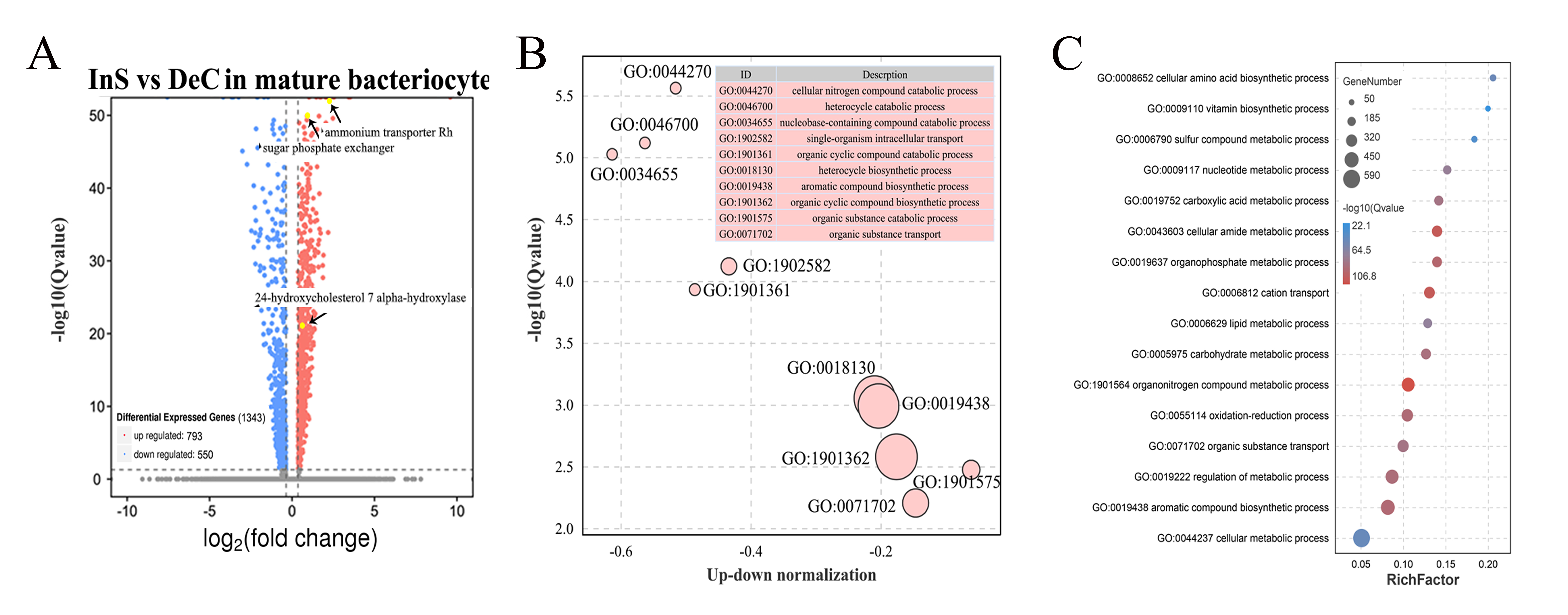

### Supplementary Fig. S9

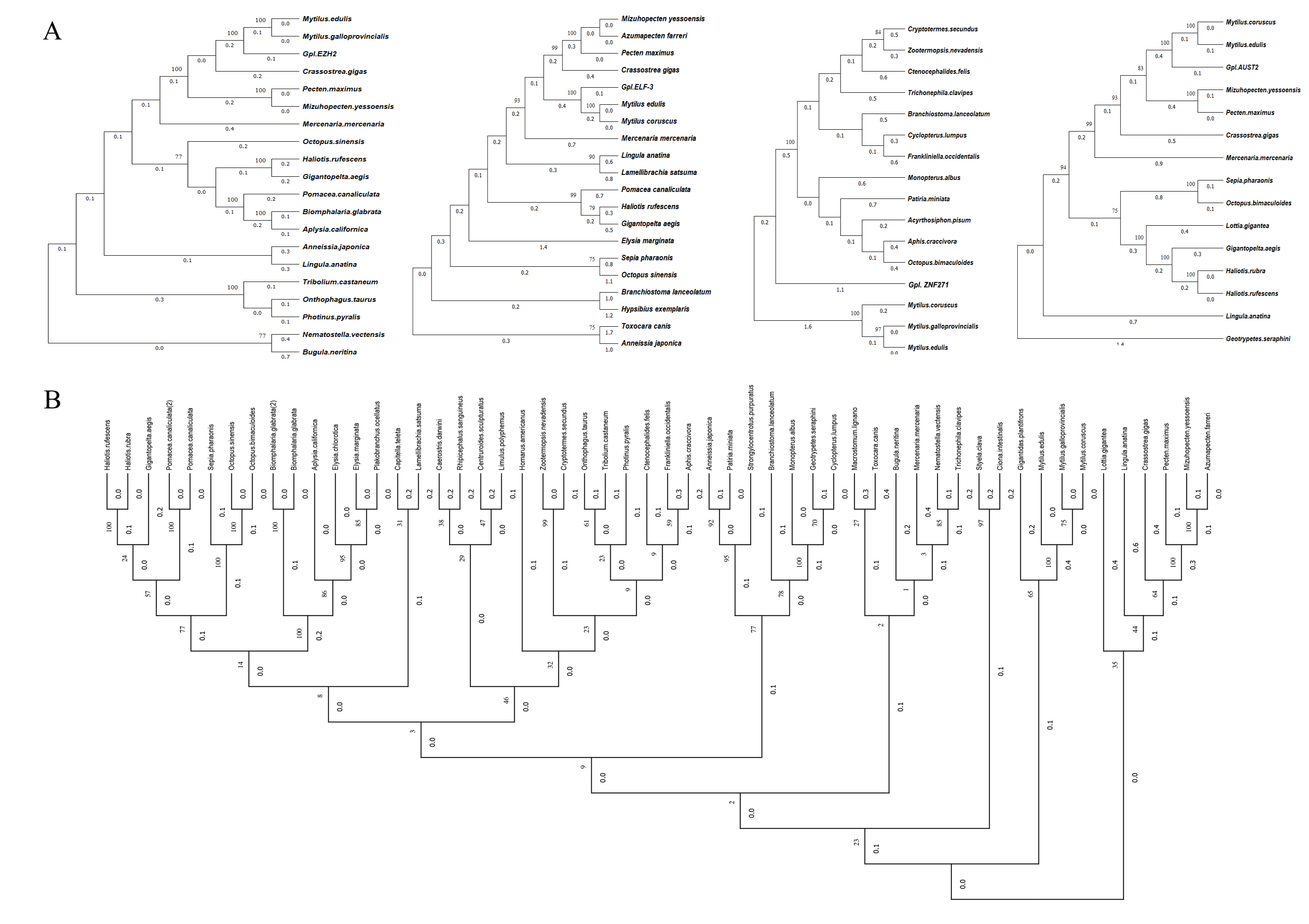
